# Allosteric Modulation of Conformational Dynamics in Human Hsp90α: A Computational Study

**DOI:** 10.1101/198341

**Authors:** David L. Penkler, Canan Atilgan, Özlem Tastan Bishop

## Abstract

Central to Hsp90’s biological function is its ability to interconvert between various conformational states. Drug targeting of Hsp90’s regulatory mechanisms, including its modulation by co-chaperone association, presents as an attractive therapeutic strategy for Hsp90 associated pathologies. Here, we utilize homology modeling techniques to calculate full-length structures of human Hsp90α in closed and partially-open conformations. Atomistic simulations of these structures demonstrated that bound ATP stabilizes the dimer by ‘tensing’ each protomer, while ADP and apo configurations ‘relax’ the complex by increasing global flexibility. Dynamic residue network analysis revealed regions of the protein involved in intra-protein communication, and identified several overlapping key communication hubs that correlate with known functional sites. Perturbation response scanning analysis identified several potential residue sites capable of modulating conformational change in favour of interstate conversion. For the ATP-bound open conformation, these sites were found to overlap with known Aha1 and client binding sites, demonstrating how naturally occurring forces associated with co-factor binding could allosterically modulate conformational dynamics.

## Introduction

The 90 KDa heat shock protein (Hsp90) is a highly conserved molecular chaperone that plays a central role in maintaining cellular homeostasis in organisms of all kingdoms of life with the exception of the archea (Taipale *et al*, 2010). Hsp90’s biological function is centered around is ability to facilitate the folding, maturation, and trafficking of a wide array of client peptides (Young *et al*, 2001; McClellan *et al*, 2007; Zhao *et al*, 2005). The related functions of these clients implicate Hsp90 with numerous cellular functions, ranging from signal transduction to complex regulatory mechanisms, as well as innate and adaptive immunity (Pearl & Prodromou, 2006). Hsp90 is thus uniquely positioned at the crossroads of several fundamental cellular pathways, representing a central hub in protein interaction networks (Taipale *et al*, 2010). Also included in Hsp90’s broad client base are known disease related peptides, association of which implicate the chaperone in the progression of several pathologies including various protein misfolding disorders, cancer, and neurological diseases (Schopf *et al*, 2017). Indeed, it has become increasingly clear that deregulation of Hsp90 may present an attractive strategy for disease treatment, leading to a growing interest in Hsp90 as a viable drug target particularly in the field of molecular oncology (Whitesell & Lindquist, 2005; Jhaveri *et al*, 2012; Taldone *et al*, 2014) and parasitology (Pallavi *et al*, 2010; Roy *et al*, 2012; Faya *et al*, 2015).

Biochemical and structural studies have revealed that Hsp90 exists as a homodimer, and that each protomer is characterized by three well-defined domains (Wayne & Bolon, 2007; Mayer & Le Breton, 2015; Schopf *et al*, 2017): the N-terminal domain (NTD) is responsible for ATP binding and hydrolysis as well as facilitating protomer dimerization (Prodromou *et al*, 1997), the middle domain (M-domain) contributes to ATPase activation (Meyer *et al*, 2003), and provides a large surface area suitable for Hsp90’s broad range of client protein interactions, and the C-terminal domain (CTD) provides the primary inter-protomer dimerization site (Ali *et al*, 2006; Harris *et al*, 2004). The NTD and M-domain are connected via a highly flexible charged linker that is thought to be involved in modulating chaperone function (Hainzl *et al*, 2009; Tsutsumi *et al*, 2009, 2012; Jahn *et al*, 2014). Hsp90’s ability to bind and release client proteins revolves around a complex nucleotide dependent conformational cycle in which the dimer transitions between a catalytically inactive open state and an active closed state (**Figure 1**). Crystal structures of the *Escherichia coli* Hsp90 homolog, HtpG, revealed that in the nucleotide free state, dimerization occurs at the CTD alone and Hsp90 adopts an open “v-like” conformation in which the M-domains are suitably exposed in preparation for client loading (Shiau *et al*, 2006; Dollins *et al*, 2007; Southworth & Agard, 2011) (**Figure 1** – A). Binding of ATP triggers conformational rearrangements that lead to NTD dimerization and an eventual conformational transition toward the closed catalytically active state. Central to this conformational switch is the behaviour of the ATP-lid which must close over the nucleotide binding pocket to entrap the bound nucleotide, and in doing so exposes essential hydrophobic surfaces which facilitate NTD dimerization (**Figure 1** – B) (Ali *et al*, 2006; Prodromou, 2012). Transition to the closed state and full ATPase activation is an inherently slow process, recording time constants in the order of minutes (Prodromou *et al*, 1997; Scheibel *et al*, 1997; McLaughlin *et al*, 2002). The transition is likely to occur via structural intermediates, which may present further energy barriers that must be overcome (Hellenkamp *et al*, 2016; Schulze *et al*, 2016). Crystal structures of mammalian Hsp90, Grp94, have revealed structurally identical partially open/closed putative intermediate conformations of Hsp90 when bound by either ADP or non-hydrolysable ATP (Dollins *et al*, 2007) (**Figure 1** – B and D). In addition to nucleotide regulation, co-chaperone interactions have been shown to play an important role in the modulation of Hsp90’s ATPase activity (Prodromou, 2012; Röhl *et al*, 2013). Interaction with the co-chaperone Aha1 has been shown to increase ATPase activity 10-fold, by promoting the closing transition through the facilitation of ATP-lid closure (Schulze *et al*, 2016) and structurally reorienting an essential M-domain catalytic loop towards the ATPase site, where it is required to co-ordinate with the γ-phosphate of ATP (Meyer *et al*, 2003). Hsp90’s catalytically active conformation (**Figure 1** – C) has been well described through crystal structures of yeast Hsp90 (Ali *et al*, 2006) and human Hsp90_ß_ (Verba *et al*, 2016). Both structures were crystalized in complex with ATP, and demonstrate structurally similar closed conformations in which the protomers are partially twisted around one another, with fully closed ATP-lids, and projection of the catalytic loops into the nucleotide binding pockets. This closed conformation is thought to be stabilized by NTD interactions with the co-chaperone P23/sba1 (Ali *et al*, 2006) (**Figure 1**-C). Eventual ATP hydrolysis and the subsequent binding of ADP triggers a conformational shift back to the open state. Under these conditions, Aha1 and P23 / sba1 are released permitting protomer uncoupling at the NTD. The subsequent repositioning of the ATP-lid back to its open conformation (**Figure 1**-D) allows for ADP release, completing the conformational switch back to the nucleotide free open conformation (**Figure 1**-A).

**Figure 1:**
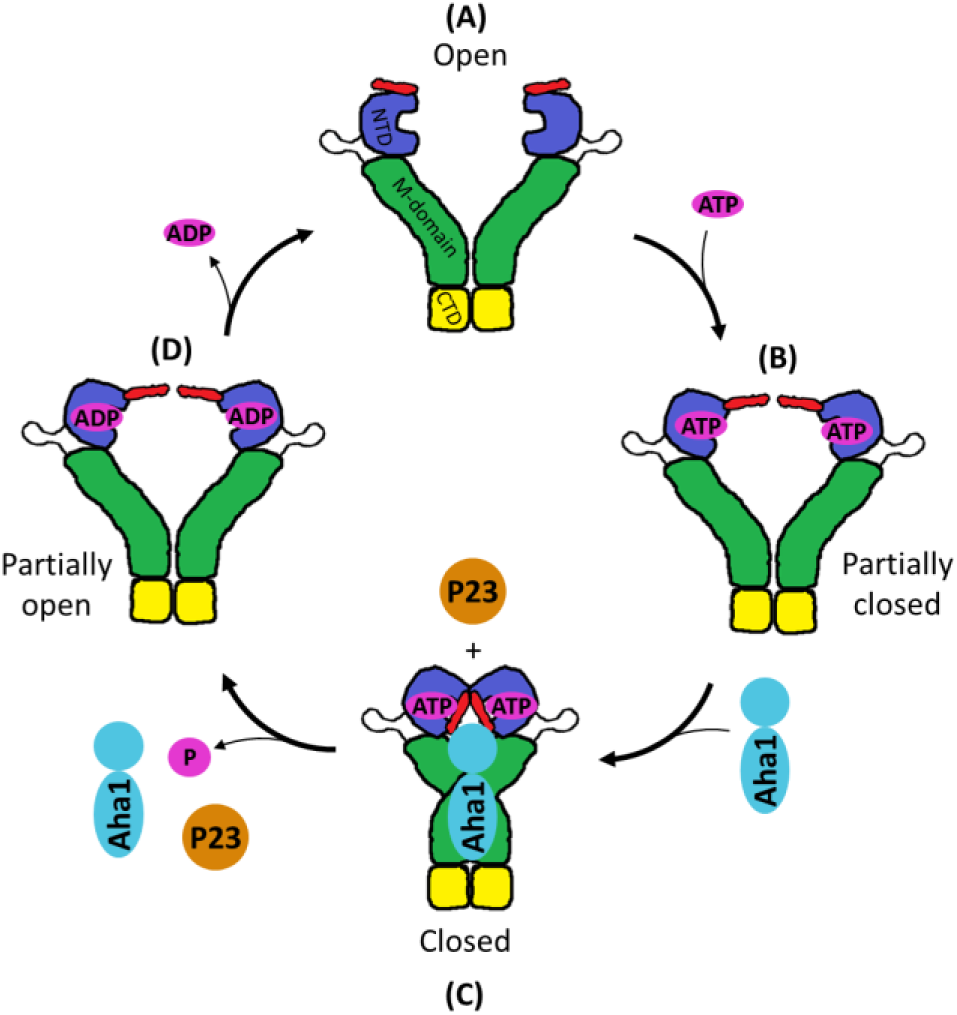
Hsp90’s nucleotide dependant conformational cycle. (A) Nucleotide free open conformation; (B) ATP binding induces NTD restructuring and the slow closure of the ATP-lid; (C) interactions with the co-chaperone Aha1 accelerates ATP-lid closure and promotes NTD dimerization, completing the conformational transition towards the closed state; (D) ATP hydrolysis and subsequent binding of ADP triggers protomer uncoupling and a transition toward the open conformation. Release of ADP returns the dimer to its original state. Domains are colored NTD (blue); M-domain (green); and CTD (yellow). The ATP-lid is shown in red.

To date, a number of computational studies aimed at elucidating the mechanistic regulation of Hsp90’s conformational dynamics have been carried out based largely on the available experimental structural data. Atomic resolution models of allosteric regulation in Hsp90 have been proposed, in which reported cross-talk between the NTD and CTD implicated ATP binding and hydrolysis at the NTD to have a strong influence on conformational dynamics at the CTD (Colombo *et al*, 2008; Morra *et al*, 2009). These studies lead to the prediction of allosteric inhibitor ligands that bind an allosteric target site identified in the CTD (Verkhivker *et al*, 2009; Morra *et al*, 2010; Vettoretti *et al*, 2016). Coarse grained and atomistic approaches coupled with energy landscape analysis were used to demonstrate the global motions and functional dynamics of Hsp90, reporting a network of conserved regions within the protein that may be important for regulating intra-protomer communications in response to ATP hydrolysis (Dixit & Verkhivker, 2012). A comprehensive all atom simulation study of several Hsp90 crystal structures from different organisms demonstrated how the functional dynamics of Hsp90 are modulated by the motion of quasi-rigid domains involving two sets of conserved residues that form inter-domain hinges between the NTD / M-domain and M-domain / CTD (Morra *et al*, 2012). Atomistic simulations coupled with force distribution analysis of the HtpG structure revealed allosteric communication pathways connecting a similar M-domain hinge and putative client binding site with the NTD nucleotide binding site (Seifert *et al*, 2012). Thermodynamic analyses of molecular dynamics (MD) simulations were used to describe structural rearrangements involved in Hsp90’s conformational transitions, identifying several key interactions that may govern conformational change (Simunovic & Voth, 2012). Lastly, structural stability analysis and protein network modeling based on atomistic simulations of Hsp90 were used to characterize the evolution of interaction networks and allosteric communication during interstate transitions, reporting a small number of conserved functional residues that may be central modulators of several functions including allosteric communication, structural stability, and co-chaperone binding (Blacklock *et al*, 2014a).

It has become increasingly clear that Hsp90’s highly complex regulatory mechanisms are, to a large extent, enabled by its interaction with a broad range of co-chaperones that are thought to tailor the flexibility of the dimer to suite its functional needs. In this manner, co-chaperones could modulate the ATPase cycle of Hsp90 by controlling the conformational transitions involved in client loading and release. Functional dynamics investigations of Hsp90’s interactions with the co-chaperones Aha1 and p23 demonstrated that binding of these cofactors alloserically modulate Hsp90’s conformational dynamics (Blacklock *et al*, 2013). Furthermore, analysis of the interaction networks involving Hsp90 complexes with the cofactors Cdc37, Sgt1, Rar1 (Blacklock *et al*, 2014b), and P53 (Blacklock & Verkhivker, 2013) demonstrated how small-world networks involving highly connected residues at the inter-protein interfaces correspond to known functional sites. Thus far, therapeutic efforts have focused largely on targeting the nucleotide binding site. However, with the implication of co-chaperone interactions for client based function, drug discovery efforts are moving towards the targeting of specific co-chaperone binding sites, or allosteric pockets, with the aim of disrupting Hsp90’s conformational dynamics and thus its loading and release of selective clients. Interestingly, the associated allosteric regulation of Hsp90 by co-chaperone interactions is believed to only occur in the cytosolic form of the protein (Johnson, 2012; Schopf *et al*, 2017) which would further minimize the selectivity of potential inhibitors. Progress of these therapeutic studies is limited however, by the lack of suitable structures of human Hsp90α.

In this study, we integrate extensive all-atom MD simulations with dynamic residue network (DRN) and perturbation response scanning (PRS) analysis techniques, to probe intra-protein allostery in full length models of human Hsp90α (**Figure 2**). Homology modeling techniques were employed to calculate accurate 3D structural models of human cytosolic Hsp90_α_ in conformations representative of the fully closed and partially-open intermediate states. By alternating the NTD configuration of each, nucleotide bound (ATP/ADP) and apo forms of the protein were prepared as suitable starting structures for all-atom MD simulations. Analysis of the global dynamics revealed that nucleotides have a minimal effect on the conformational change in the closed complexes, but that bound ADP in the partially open (PO) structure results in significant conformational rearrangements representative of the fully-open (FO) conformation and thus the opening transition. Analysis of the internal dynamics revealed that bound ATP ‘tenses’ and structurally stabilizes the dimer regardless of its conformation, while the ADP and apo configurations ‘relax’ the protein, imparting conformational flexibility in the protomers. DRN analysis revealed several nucleotide independent communication hubs common in both conformations which overlap with known functional sites. PRS analysis confirmed these findings by accentuating many of these same sites, and revealed several key allosteric effector residues that may be involved in modulating conformational dynamics. Included in these are residues that have been previously implicated in Aha1 binding interactions, demonstrating how the external force perturbations utilized in the PRS analysis could be used to identify naturally occurring perturbations arriving through co-chaperone binding. Collectively, the data presented here provides a suitable platform for novel experimental investigations regarding human Hsp90α chaperone function and regulation.

**Figure 2:**
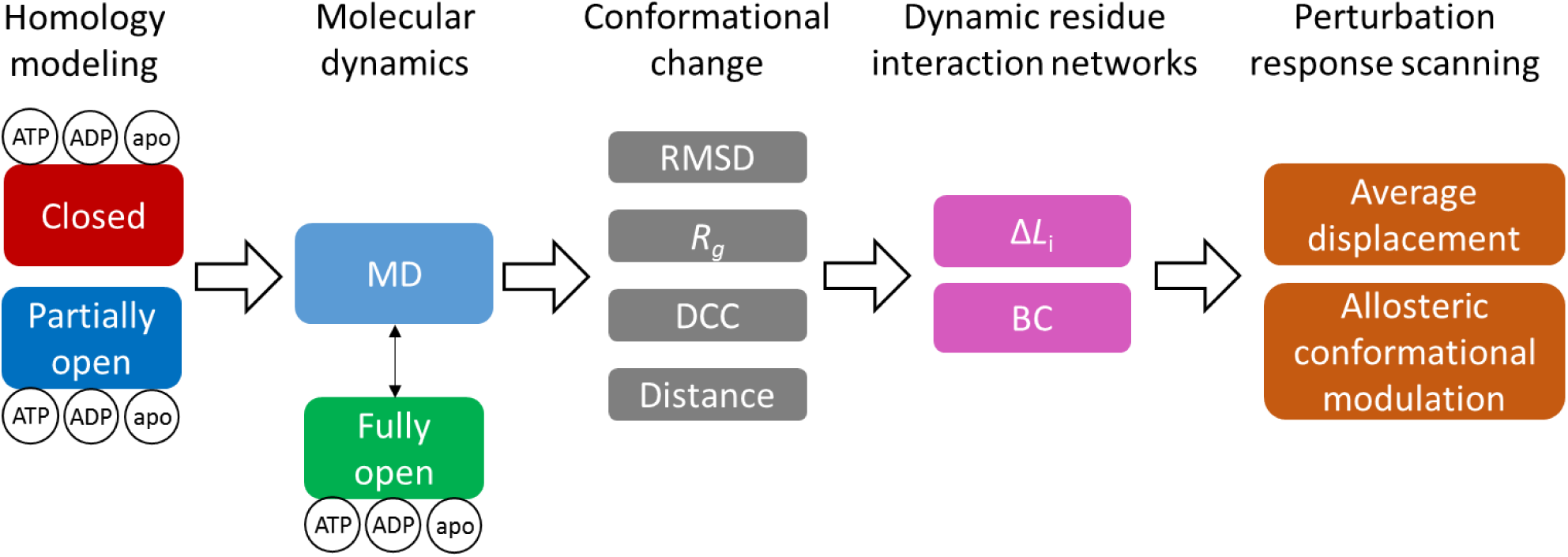
Schematic overview of methodology. The Fully open structure was obtained from MD simulation of the partially open complex

## Methodology

### Homology modeling

The 3D structure of human cytosolic Hsp90α was modeled in an open and closed conformation using the full protein sequence NP_005339.3 as the target sequence by the MODELLER (Šali, 2013) program. For the closed conformation, the full length cryo-EM structure of human Hsp90β in complex with ATP (PDB id: 5FWK) (Verba *et al*, 2016) was used as a modeling template. 5FWK represents the full protein structure except for the linker between the NTD and MD (res 228-277), which due to its inherent flexibility is yet to be crystallised. For this reason, residues 230-275 were deleted from the target sequence, and replaced with four glycine residues. Modeling of the open conformation required the use of a multi-template modeling approach to ensure a ‘v-like’ open dimer, with an extended orientation for each protomer. The canine endoplasmic reticulum paralog of Hsp90, GRP94, demonstrates a partially open ‘v-like’ conformation with dimerization at the CTD (PDB id: 2O1U) (Dollins *et al*, 2007), and was used as the primary template. Given that 2O1U and the target sequence only share 53% sequence identity, 5FWK was used as a secondary template, providing cover for the missing region L290-R299. 2O1U lacks structural data for the ATP-lid domain (res 98-123), while 5FWK is representative of the structure with a closed ATP-lid. Thus, to ensure that the ATP-lid was modeled in its open conformation, the NTD (res 15-278) of 5FWK were excluded from the alignment file and the open ATP-lid NTD crystal structure of human Hsp90 (PDB 3T10) was used as a second template for this domain. A total of 100 models were built for each dimerized conformation (open and closed), and the respective models with the lowest normalized DOPE score (DOPE-Z score) selected for further evaluation by VERIFY3D (Eisenberg *et al*, 1997), Errat (Colovos & Yeates, 1993) and ProSA (Wiederstein & Sippl, 2007). To obtain each of these conformations in complex with ATP and ADP, the respective nucleotide’s structural coordinates were extrapolated from representative crystals structures by superimposing PDBs 3T0Z (Li *et al*, 2012) and 1BYQ (Obermann *et al*, 1998) for ATP and ADP respectively. The apo NTD was obtained by omitting nucleotide coordinate data. In this manner a total of six human Hsp90α configurations were prepared, three in the open conformation and three in the closed conformation.

### Molecular dynamics simulations

MD simulations were carried out on all nucleotide bound configurations of the homology models. All MD simulations were performed using GROMACS 5.1.2 (Berendsen *et al*, 1995) with in the CHARMM 36 force field (MacKerell *et al*, 1998; Mackerell *et al*, 2004; Best *et al*, 2012), using a orthorhombic periodic box with a clearance space of 1.5 nm. Water molecules were added as solvent, and modeled with the TIP3P water model. The system was neutralized using a 0.15 M NaCl concentration. Prior to production runs, all molecular systems were first energy minimized using a conjugate-gradient being energy relaxed with up to 50 000 steps of steepest-descent energy minimization, and terminated when the maximum force < 1000.0 kJ/mol/nm. Energy minimization was followed by equilibration, first in the *NVT* ensemble at 310 K using Berendsen temperature coupling, and then in the *NPT* ensemble at 1 atm and 310 K until the desired average pressure (1 atm) was maintained and volumetric fluctuations stabilized. All production simulations were run for a minimum of 100 and maximum of 200 ns, to ensure that the backbone root-mean-square deviation (RMSD) of the protein equilibrated with a fluctuation of no more than 3 Å for at least 20 ns. Data pertaining to the protein and nucleotide were saved at 2 ps intervals for further analysis. All simulations utilized the LINCS algorithm for bond length constraints. The fast particle mesh Ewald method was used for long-range electrostatic forces, the switching function for van der Waals interactions was set to 1.0 nm and the cutoff set to 1.2 nm. NH3^+^ and COO^−^ groups were used to achieve the zwitterionic form of the protein. Periodic boundary conditions were applied in all directions and the simulation time step was set to 2 fs.

### Dynamic cross-correlation

Dynamic cross-correlation (DCC) describes the correlation of motions between each C_α_ atom in a protein over the course of an MD trajectory. DCC is calculated based on the reduction and normalization of the 3*N* x 3*N* covariance matrix (**C**). The MD-TASK (Brown *et al*, 2017a) software suite was used to calculate DCC for each MD trajectory. **C** is constructed from the MD trajectory sampling data at 100 ps intervals, and the average deviation of each C_α_ atom from a mean structure representative of the trajectory length is calculated. The essential directions of correlated motions are determined by diagonalizing **C** to get the *N* x *N* correlation matrix (**Corr**):

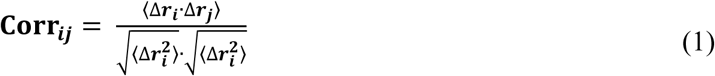

### Dynamic residue network analysis

To analyse inter-and intra-domain communication, the protein is represented as a residue interaction network (RIN), where the C_β_ atoms of each residue (C_α_ for glycine) are treated as nodes within the network, and edges between nodes defined within a distance cut off of 6.7 Å (Atilgan *et al*, 2004). In this study, the MD-TASK (Brown *et al*, 2017a) software suite is used to construct dynamic RINs (DRN) based on the all-atom MD trajectories. Each DRN is the combination of RINs representative of time points (frames) in the MD trajectory, using a 200 ps time interval between frames. MD-TASK was also used to analyse each DRN in terms of change in reachability (Δ*L*_*i*_) and betweenness centrality (BC) for each residue. MD-TASK calculates *L* and BC matrices for each RIN in the DRN, and calculates Δ*L*_*i*_ by finding the difference between every *n*^th^ RIN and the initial RIN at 0 ps. Finally, MD-TASK averages the Δ*L*_*i*_ and BC matrices. Average Δ*L*_*i*_ and BC indicates residues that experience permanent changes in Δ*L*_*i*_ and BC as opposed to minor fluctuations over the course of the MD trajectory (Brown *et al*, 2017a). For comparative purposes, BC values are normalized in the range 0.0 – 1.0.

### Perturbation response scanning

Perturbation response scanning (PRS) is a computational technique used to predict the relative response of all residues in a protein, to an external force perturbation, such as ligand binding, at a single residue. The theory of PRS has been thoroughly described in previous studies (Atilgan & Atilgan, 2009; Atilgan *et al*, 2010; Penkler *et al*, 2017); the algorithm is available in the MD-TASK software suite (Brown *et al*, 2017a). In brief, PRS utilizes linear response theory (LRT) to approximate the shift in coordinates (**R**_1_) of a given protein conformation (**R**_0_) in response to a perturbation of the Hamiltonian (Yilmaz & Atilgan, 2000; Ikeguchi *et al*, 2005; Atilgan & Atilgan, 2009).

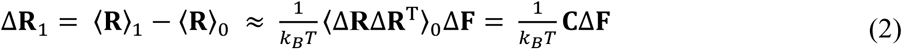

The subscripts 0 and 1 represent the unperturbed and perturbed protein structures, respectively. The force vector **F** describes the coordinates of the inserted force perturbation on a select reside. **C** is the covariance matrix, which MD-TASK constructs directly from an equilibrated portion of the MD trajectory (20 ns in this work). For each PRS experiment, the initial and final structures (states 0 and 1) were defined as follows: The first frame of the equilibrated MD trajectory segment was set as state 0. For the open to closed transitions, the final structure (state 1) was set to the equilibrated structure of the ATP bound closed configuration. Conversely, for the closed to open transitions, the final structure was set to the equilibrated ADP bound open conformation after 200 ns. Each residue in the initial state was sequentially perturbed with 250 fictitious forces of random direction. For each residue *i*, MD-TASK assesses the quality of the response vector (*ΔR*_*k*_) using the Pearson’s correlation coefficient by correlating Δ*R*_*k*_ with the experimental displacements averaged over all affected residues *k* using the following equation:

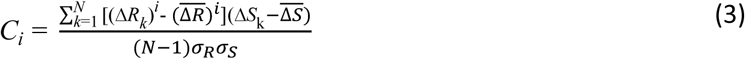

where a *C*_*i*_ value close to 1 implies a good agreement with the experimental change and a *C*_*i*_ value close to 0 indicates little or no agreement. In each case only the maximum *C*_*i*_ is recorded for each residue. We have previously discussed and shown the reproducibility of this approach with large multi-domain proteins with complex rotational and translational conformational restructuring (Penkler *et al*, 2017).

## Results and Discussion

### Homology modeling

Structures of human Hsp90α in closed and partially open ‘v-like’ conformations were obtained by homology modeling (**Figure 3**). In each case 100 unique homology models were calculated and ranked by normalized DOPE-Z score. The normalized DOPE Z-score is an atomic distance-displacement statistical potential used to evaluate the fold of a model, and is dependent on a sample structure of a native protein (Shen & Sali, 2006; Eramian *et al*, 2008). A value less than-0.5 is deemed an accurate representation of the native structure (Eswar *et al*, 2001), and the lowest scoring models were selected for further model evaluation and verification. Table **SD 1** summarizes the results of the best scoring models after separate model evaluation with Verify3D, Errat, and ProSA techniques. Verify3D is an online web-server that determines the relative compatibility of an atomic coordinate structure (3D) with its own amino acid sequence (1D), assigning structural classes based on secondary structure, location, and local environment, comparing the results with known good structures (Bowie *et al*, 1991; Luethy *et al*, 1992). The percentage of residues within the protein that score more than 0.2 is reported, and a minimum of 80% is required to pass model evaluation. Errat is a program that analyzes the statistics of all non-bonded interactions between carbon, nitrogen and oxygen atoms reporting the percentage of residues that fall below a rejection confidence level of 95% (Colovos & Yeates, 1993). The ProSA web-server calculates the overall quality of a protein structure based on a Z-score which indicates the goodness of fit between the modeled structure and native structures of a similar size (Wiederstein & Sippl, 2007). Both, the selected open and closed homology models passed each of these evaluation tests, recording scores in the good to very good range (**SD 1**). Model evaluation was carried out before and after energy minimization, to assess the impact of possible bad contacts with in the structure. The erroneous regions highlighted by these evaluation techniques pre-minimization were all confined to loop regions, particularly the missing linker region between the NTD and the M-domain. The quality of these regions improved after energy minimization, the final models providing suitable starting structures for long range MD simulations.

**Figure 3:**
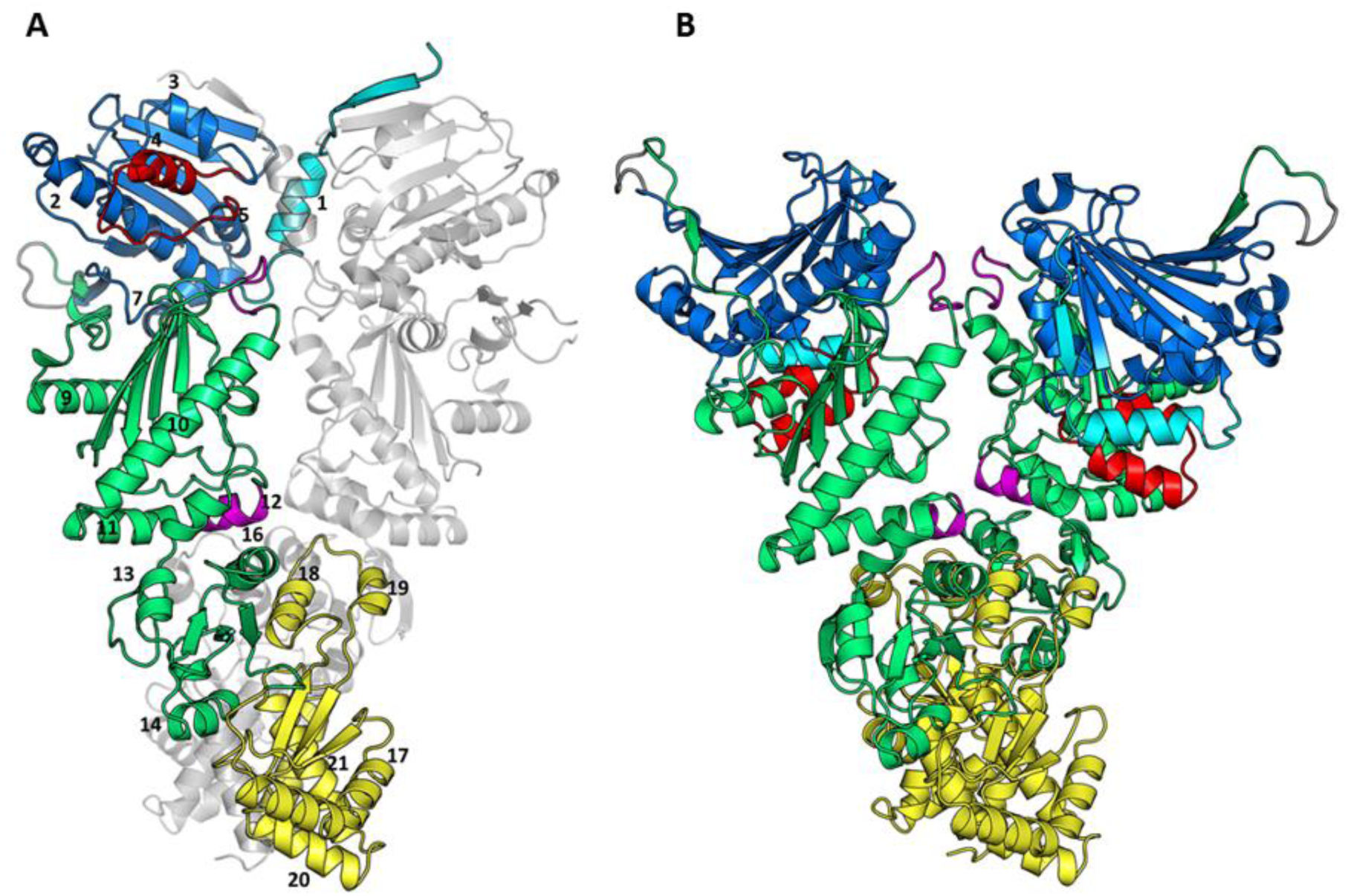
3D Illustration of the human Hsp90α homology models. (A) Fully closed conformation, (B) partially open conformation. Indicating the locations for the NTD (blue); charged linker (grey); M-domain (green); CTD (yellow); ß-strand_1_-helix_1_ interface (cyan); ATP-lid (red); and M-domain/CTD hinge (magenta). Helices are labelled according to the numbering convention used throughout the text.

### Effect of nucleotide binding on conformational change

The NTD nucleotide bound configuration (ATP / ADP / apo) has been intricately coupled with the modulation of conformational dynamics in Hsp90’s chaperone cycle (**Figure 1**). To determine the relative effect of bound nucleotide on the conformational dynamics (opening and closing transitions) of human Hsp90, all-atom MD simulations in explicit water were carried out on nucleotide bound (ATP / ADP) and apo closed, PO, and FO complexes, in a total of nine unique simulation runs (**Table 1**). The closed and PO simulations were initiated based on the homology models, while the FO simulations were based on a structure extracted from the PO-ADP MD trajectory. The resultant trajectories were analysed in terms of their backbone RMSD, residue root-mean-square-fluctuations (RMSF), radius of gyration (*R*_*g*_), inter-protomer distance, dynamic cross-correlation (DCC) of atomic motions, and observed conformational changes.

**Table 1:**
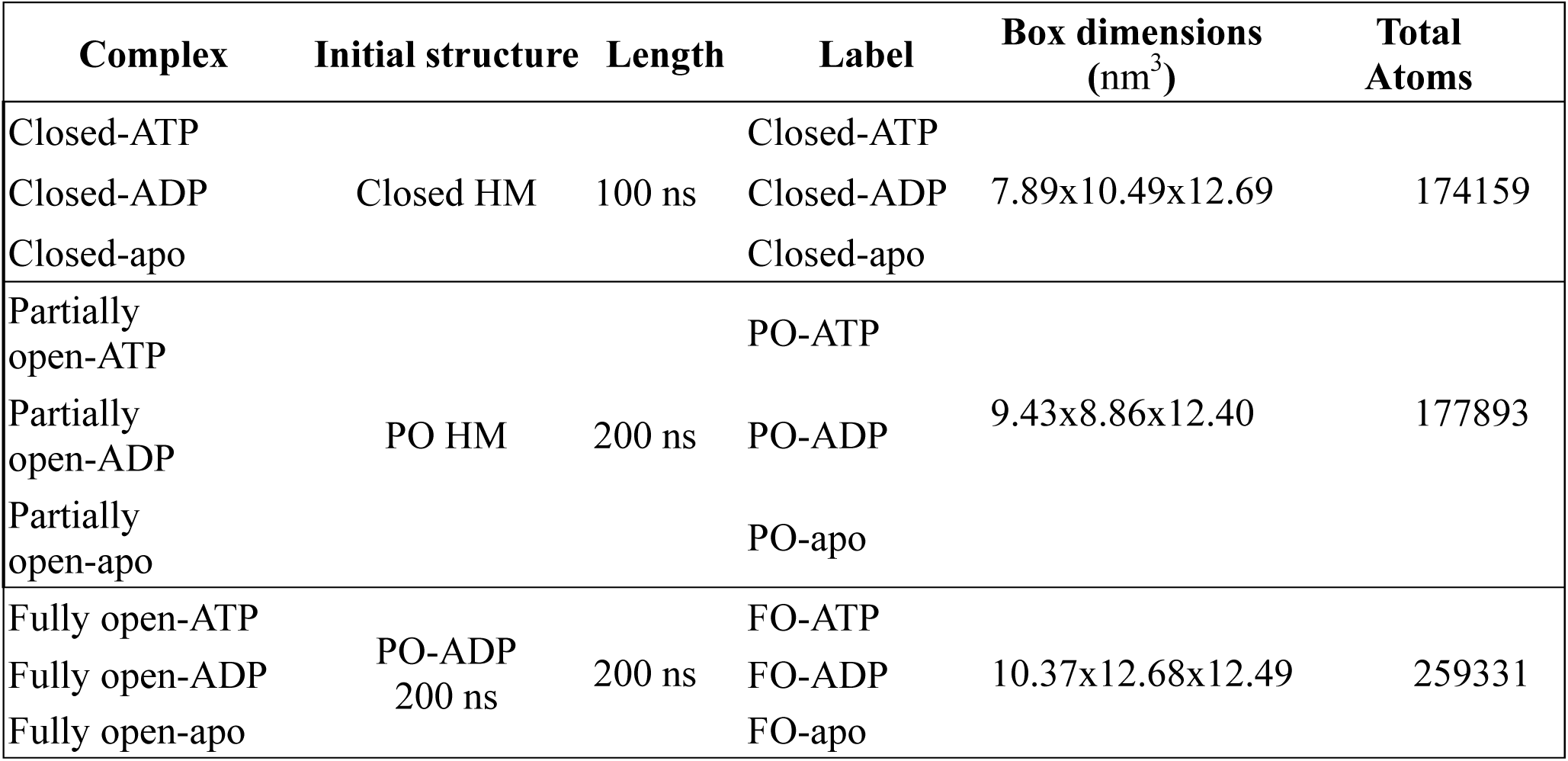
Summary of MD simulations for the closed, partially open, and fully open complexes. HM: Homology model; PO: Partially open; FO: Fully open.

| Complex | Initial structure | Length | Label | Box dimensions (nm<sup>3</sup>) | Total Atoms |
| --- | --- | --- | --- | --- | --- |
| Closed-ATP | Closed HM | 100 ns | Closed-ATP | 7.89x10.49x12.69 | 174159 |
| Closed-ADP |  |  | Closed-ADP |  |  |
| Closed-apo |  |  | Closed-apo |  |  |
| Partially open-ATP | PO HM | 200 ns | PO-ATP | 9.43x8.86x12.40 | 177893 |
| Partially open-ADP |  |  | PO-ADP |  |  |
| Partially open-apo |  |  | PO-apo |  |  |
| Fully open-ATP | PO-ADP<br>200 ns | 200 ns | FO-ATP | 10.37x12.68x12.49 | 259331 |
| Fully open-ADP |  |  | FO-ADP |  |  |
| Fully open-apo |  |  | FO-apo |  |  |

#### Closed complexes

Backbone RMSD analysis of the closed complexes indicates fairly stable structures for all three configurations over 100 ns trajectories, with RMSD converging around 5-6 Å (**Figure 4**-A). The structures at the end of 100 ns simulations are displayed in **SD 2.** RMSF analysis (**SD 3**) reveals large fluctuations of loop R60-E75, the ATP-lid (res 112-136), and the charged linker (res 220-282). Despite similar RMSD convergence and RMSF values between the three configurations, analysis of their final structures (100 ns) reveals subtle differences in the NTD of each complex when compared to their initial structures (0 ns) (**SD 2**). The NTD of the ATP complex appears to become more compact compared to the NTDs of ADP and apo complexes. Analysis of *R*_*g*_ over the MD trajectory provides a metric for evaluating the degree of compactness of a protein of interest (Lobanov *et al*, 2008). *R*_*g*_ for the closed complexes shows an initial decrease for the ATP complex which eventually stabilizes around 36.5 Å (**Figure 4**-B), suggesting a tightening of the dimers around the central axis. The ADP and apo complexes show a smaller initial decrease compared to the ATP complex, both stabilizing around 38.5 Å, suggesting these complexed to be more ‘relaxed’ around the central axis. The apparent compactness of the ATP complex is also confirmed when analysing the MD trajectories in terms of correlated C_ß_ atom motions (1). In this analysis, each correlation matrix (**Figure 4**-C) represents the correlation between any two C_ß_ atoms over the course of an MD trajectory. Positive values approaching 1 describe correlated motions (in the same direction), and negative values close to-1 describe anti-correlated motions (in opposite directions). In the ATP complex, strong anti-correlation is observed between the NTD and CTD within and between individual protomers, suggesting these domains to have anti-correlated motions either towards or away from each other in the presence of ATP. The degree of anti-correlation between these domains is reduced in the ADP and apo complexes, especially in protomer 2, supporting the hypothesis that these complexes resemble more relaxed structures. Although the differences observed between the two protomers may be attributed to the MD simulation length, it may also be evidence of asymmetric conformational dynamics. In fact, the more detailed DRN and PRS analyses we present later in the manuscript (**Figure 2**), display how residues in the two protomers differentiate in function, in line with experimental findings towards this end (Flynn *et al*, 2015). Protomer uncoupling at the NTD precedes transition towards the open state. Looking at the motions between the NTDs of each protomer, positive correlated motions are observed between the NTDs of the nucleotide bound complexes, suggesting that the NTDs move in a concerted fashion relative to one another. Strikingly, the converse is observed for the apo structure, in which the NTDs show anti-correlated motions relative to one another, suggesting movement in opposite directions, possibly representing protomer uncoupling.

**Figure 4:**
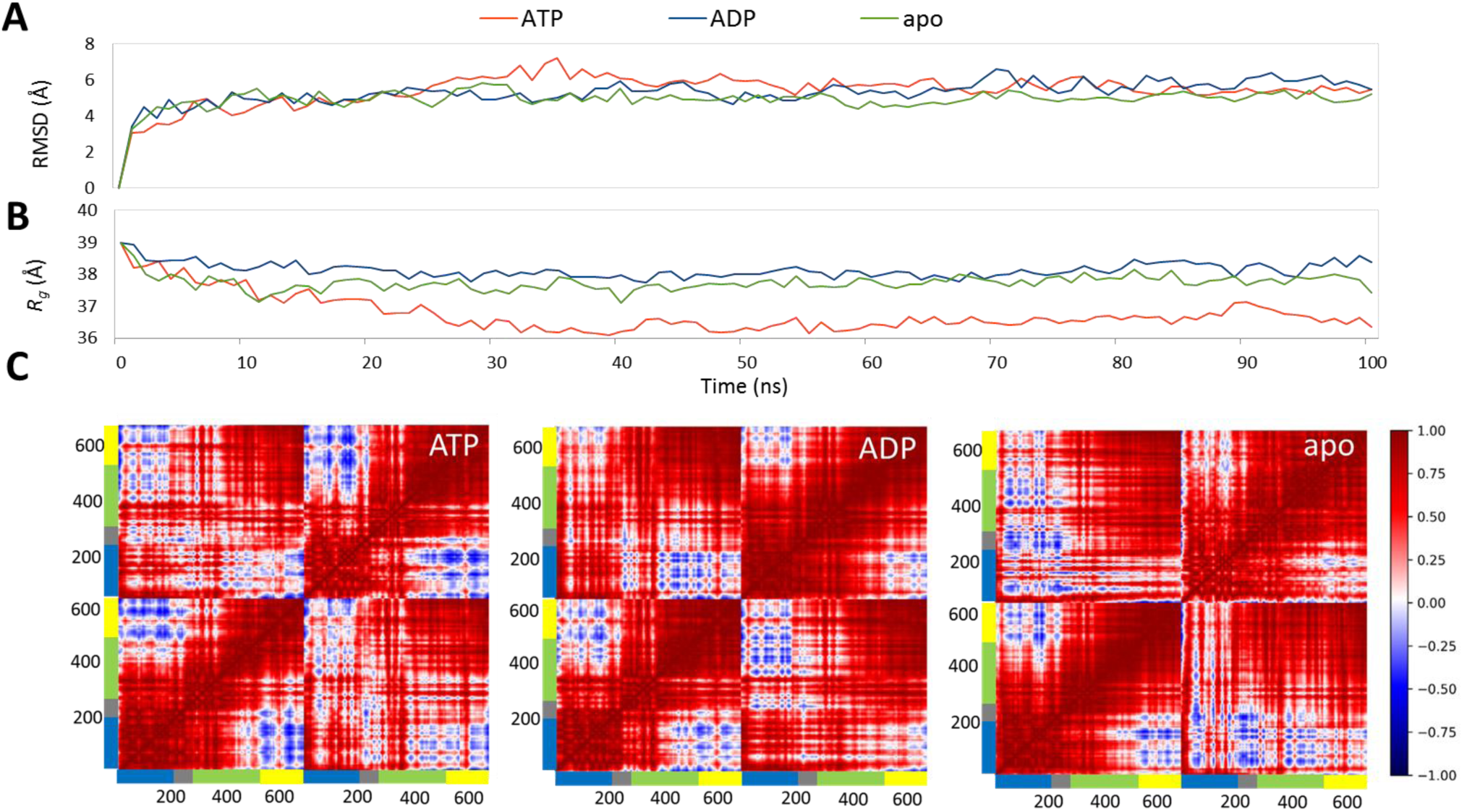
Global conformational dynamics metrics for the closed conformation complexes: (A) backbone RMSD; (B) radius of gyration; (C) DCC heat maps showing correlated motions (positive) in red and anti-correlated motions (negative) in blue. The color bar accompanying each heatmap represents each functional domain: NTD (blue), linker (grey), M-domain (green), and CTD (yellow)

Previous experimental studies have shown evidence that Hsp90’s conformational transition from the closed ATP bound ‘tense’ state to the open ‘relaxed’ state can only be achieved after ATP hydrolysis and the subsequent release of ADP from the NTD (Schopf *et al*, 2017). The MD data presented here suggests that 100 ns runs are insufficient to observe the expected global restructuring required for the opening transition, with similar final structures observed for all three complexes (**SD 2**). However, the data presented do demonstrate that the complex is amenable to subtle structural changes and deviation from the initial structure under the different NTD configurations. Bound ATP appears to enhance the NTD-CTD ‘compactness’ of the complex, providing evidence in agreement that this complex favours the closed ‘tense’ state (Ali *et al*, 2006; Krukenberg *et al*, 2011; Flynn *et al*, 2015; Schopf *et al*, 2017), while the ADP and apo complexes appear to relax the NTD and CTD relative to one another, possibly in preparation for the opening transition.

#### Partially open complexes

The PO complexes of human Hsp90α were calculated based primarily on the canine Grp94 template (PDB id: 2O1U), which is representative of a partially open structure that may represent the native conformation of the protein in the absence of bound client proteins or co-factors (Dollins *et al*, 2007). The atomic coordinates for the ATP-lid are missing from 2O1U, and as such these complexes were modeled with the ATP-lid in the open configuration, providing a putative intermediate structure between the closed catalytically active state and the native inactive fully open state.

The backbone RMSD plots of the PO complexes (**Figure 5**–A) indicate minor conformational restructuring within the first 35 ns, after which the ATP and apo complexes stabilize, converging to a value around 5 Å, while the ADP complex undergoes extensive conformational fluctuations, without equilibrating over the 200 ns simulation. Superimposing the CTDs of the final structures with the initial PO structure shows that each complex configuration yields a unique final structure after the 200 ns MD simulation (**SD 4**). As expected, the ATP complex undergoes minor conformational change in which the NTDs are reoriented towards the M-domain of the opposite protomer (**SD 4**-red). This reorientation appears to tighten the two protomers, in line with conformational rearrangements thought to be necessary for catalytic activation (Meyer *et al*, 2003; Prodromou, 2012). The ADP complex appears to undergo extensive conformational restructuring, whereby the protomers disengage from one another, in a distinct opening of the molecular clamp. Indeed, the final PO-ADP structure closely resembles an open ‘v-like’ conformation (**Figure 5**-D), the backbone RMSD deviating only 8.23 Å from the bacterial Hsp90 HtpG (PDB id: 2IOQ) (Shiau *et al*, 2006) after 3D superimposition (**SD 4**-B). Like the PO-ATP complex, the PO-apo complex also shows minor deviation from the initial structure, the only notable conformational change being the NTD of the second protomer, which appears to relax and extend in an upward flexing motion away from the M-domain of the opposite protomer (**SD 4**-green). Measuring the time evolution of NTD inter-protomer distance, defined by the distance between the center of mass of each the NTDs, over the course of each MD simulation (**Figure 5**-B) reveals that the protomers of PO-ADP complex fully uncouple after ∼20 ns, the inter-protomer distance increasing by ∼10 Å, to fluctuate around 30 Å. In contrast, both the ATP and apo complexes maintain their NTD inter-protomer distance, fluctuating around 21 Å.

**Figure 5:**
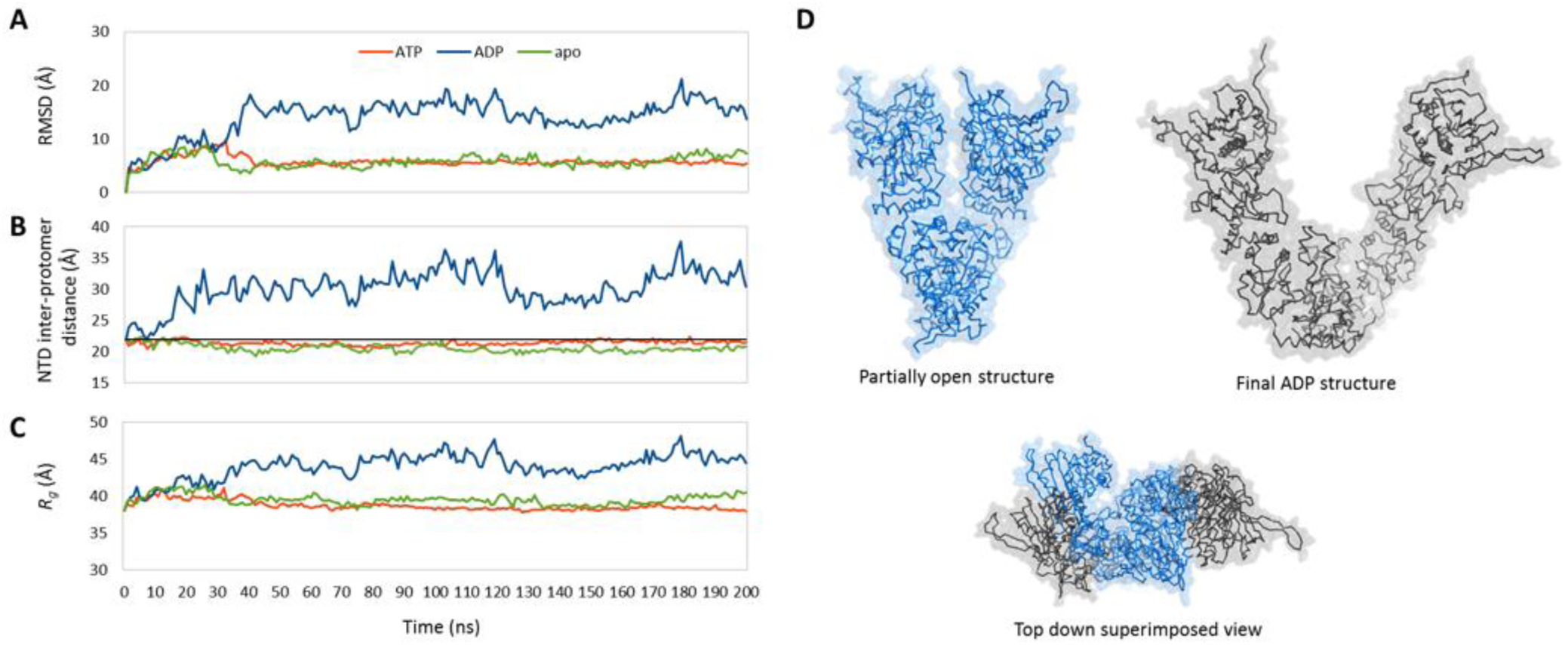
Global conformational dynamics metrics for the partially open conformation complexes: (A) Backbone RMSD, (B) NTD inter-protomer distance (black line represents the distance at 0 ns), (C) Rg plots, (D) 3D illustration of dimer opening in the PO-ADP complex, showing the initial partially open structure (blue) and the final structure after MD simulation (grey)

RMSF analysis of the PO complexes (**SD 5**) shows an elevated degree of residue fluctuation in the NTDs of all three complexes, with slightly higher fluctuations observed for the ADP complex. Notable NTD fluctuations include; the N-terminal β-strand_1_ (res 15-26), β-sheets 3-5 (res 65-75, 160-80), the ATP-lid (res 112-136), and the charged linker (res 210-290). A notable difference between the different PO configurations is observed in the M-domain (res 288-549), where fluctuations are increased for the ADP complex, particularly the catalytic loop region (res 391-406). All three complexes show similar residue fluctuations at the CTD; however, the ADP complex once again experiences higher fluctuations, this time at helix_18_ and helix_19_ (res 601-632). Collectively this data suggests that the ADP complex is more flexible than the ATP and apo complexes, experiencing fluctuations in all three subdomains which may be evidence of a ‘relaxed’ ADP state. This apparent increase in flexibility for the ADP complex may be an important structural characteristic for the destabilization of anchoring inter-protomer interactions in the M and CTDs, allowing the protomers to easily disengage as seen in the final structure of the ADP bound complex (**Figure 5**-D).The converse may be true for the PO-ATP and PO-apo complexes, in which protomer rigidity may help maintain inter-protomer interactions, thus stabilizing and favouring the closed state. *R*_*g*_ analysis for these complexes supports this observation (**Figure 5**-C), in which the PO-ADP complex becomes considerably less compact over time, increasing in value to fluctuate around 45.0 Å at approximately the same time point that the protomers first disengage (**Figure 5**-B). In contrast, the PO-ATP and PO-apo complexes maintain their original structures, fluctuating around 38.0 Å, the former becoming slightly more compact than the latter. Looking at the inter-protomer interactions present in the final MD simulation structures, the only interactions conserved in all three complexes are those involved in CTD dimerization, the notable difference being helix_18_ (res 601-632). In the initial closed and PO structures, helix_18_ is seen to form intimate interactions with the identical residues in the opposite protomer, forming an important interface between the protomers. Despite minor fluctuations of these residues over the course of the MD simulations (**SD 5**), both the apo and ATP complexes appear to retain this inter-protomer interaction, while in the ADP complex they are completely abolished, allowing helix_18_ and helix_19_ to fold back towards their respective protomers. The CTD has been previously implicated in the allosteric modulation of dimerization in yeast Hsp90 (Morra *et al*, 2009; Retzlaff *et al*, 2009; Soroka *et al*, 2012; Morra *et al*, 2010). This data could describe how ADP may allosterically modulate conformational dynamics, in which it increases protomer flexibility, allowing the NTDs to become extended triggering disengagement of helix18-19 interface, which in turn causes further destabilization of other inter-protomer interactions in the M-domains, resulting in a transition to the open state.

#### Fully open ‘v-like’ complexes

All-atom MD simulations of the PO complexes revealed drastic conformational changes in the ADP configuration that closely resembled the FO ‘v-like’ conformation (**SD 4**-B). This MD structure provided a suitable starting point for further MD investigations regarding the effect of bound nucleotide on the conformational dynamics of the FO conformation. As such, the final structure of the PO-ADP complex was selected from the 200 ns trajectory, and FO-ATP, FO-ADP, and FO-apo complexes were prepared in the same manner as the previous two conformations, and each complex submitted to 200 ns MD simulations.

The RMSD plots of the fully open complexes (**SD 6**-A) indicate minor backbone fluctuations for the ATP and apo complexes, the RMSD values converging around 6 Å. However, the FO-ADP complex appears to undergo large conformational changes at the 100 ns time point, which correspond to a slight increase in NTD inter-protomer distance (∼2 Å) (**SD 6**-B). RMSF analysis of all three complexes (**SD 7**) reveals very similar residue fluctuation profiles to the PO complexes (**SD 5**), where the NTD and M-domains experience higher fluctuations compared to the CTD. Comparing the final structures of each complex, the only noticeable difference is observed for the ATP complex in which the NTDs appear to re-orient inwards, towards their respective M-domains, a movement that may be indicative of an early closing motion.

Overall, the data presented here demonstrate that major conformational dynamics are experienced for the PO complexes in response to the ligand bound state of the NTD, while more subtle structural rearrangements are experienced by the closed and FO complexes. In each case, ATP appears to confer structural rigidity and ‘tensing’ of the dimer, while ADP enables a more ‘relaxed’ flexible complex. The apo complex engenders opposing correlated motions in the closed complexes, while it has little or no effect on the PO and FO complexes which appear largely stable. This contrasting behaviour may be in agreement with reports of stochastic dynamics observed for the apo protein (Krukenberg *et al*, 2008, 2011; Southworth *et al*, 2008). While these findings are in agreement with several previous studies (Krukenberg *et al*, 2011; Schopf *et al*, 2017; Flynn *et al*, 2015), the only conformational transition observed within the MD simulation time scales used in this study was the opening transition of the PO-ADP bound complex. To probe our data for further insights regarding nucleotide modulation of inter-state transition, the MD trajectories of the closed and FO complexes were subjected to DRN analysis to investigate the effect of bound nucleotide on intra-protein communication, as well as PRS analysis to identify single residues involved in modulating interstate conversion.

### Dynamic residue network analysis

In this analysis, each Hsp90 complex is treated as a network of residues, where the C_β_ (C_α_ for Gly) atoms for each residue are considered the nodes in the network, and the edges between nodes established based on a separation cut-off distance of 6.7 Å (Atilgan *et al*, 2004). In this manner, DRNs are calculated for each trajectory, and the resultant matrices analysed in terms of the shortest path (*L*) and betweenness centrality (BC). Briefly *L*_ij_ is the number of connections (edges) utilized to reach node/residue *i* from *j*, using the shortest possible path. *L*_*i*_ is thus the average path length to node *i*, and is calculated as the average number of steps that the node may be reached from all other nodes of the network. Here, we monitor the change in *L*_i_ (Δ*L*_i_) in response to the ligand bound configuration of the NTD, and interpret large Δ*L*_i_ as sites on the protein that may be important for inducing intra-protein communication (Ozbaykal *et al*, 2015). BC is associated with *L* in that it is proportional to the number of shortest paths passing through a given residue, thus providing a measure of the usage frequency of a residue in cross network communication, where high usage residues are considered to potentially play a role in controlling intra-protein communication (Ozbaykal *et al*, 2015). The efficiency of this method for the calculation and analysis of BC and Δ*L*_i_ along the MD simulation has previously been shown in a recent study on single nucleotide variants in the renin-angiotensinogen complex (Brown *et al*, 2017b). Here, we apply this analysis technique to each of the closed and FO complex configurations.

#### Nucleotides modulate long-range residue reachability

The Δ*L*_i_ profiles for the closed complexes (**Figure 6**), show that the ATP and apo complexes experience mostly negative Δ*L*_i_ values over the length of the protein, while the ADP complex experiences both positive and negative change. Negative Δ*L*_i_ can be described as a shortening of the average path length between all residues and residue *i*, while positive Δ*L*_i_ values indicate increased path lengths. Thus, the negative Δ*L*_i_ observed in the ATP and apo complexes may suggest these conditions favour more compact structures, resulting in decreased path lengths. Conversely, the increased path lengths observed for the ADP complex may suggest a more relaxed elongated structure. Indeed, this explanation of the findings is in agreement with our conformational dynamics results, confirming structural tensing and relaxing under ATP and ADP conditions, respectively.

**Figure 6:**
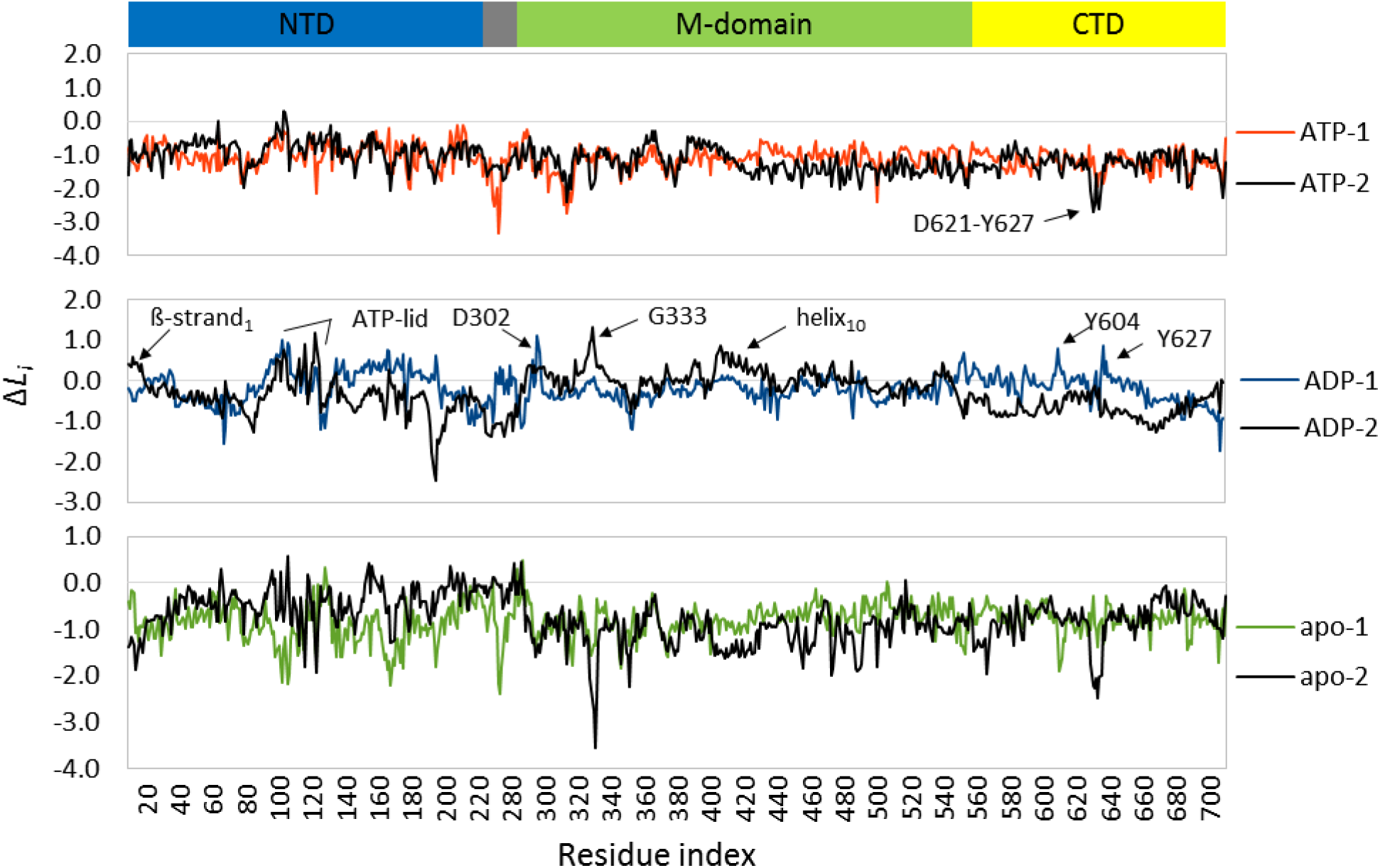
Change in reachability (*ΔL*_*i*_) plots for the closed conformation complexes. Protomer 1 is coloured by NTD configuration (ATP – red, ADP – blue, apo – green), and protomer 2 is in black.

Regions of the protein that display elevated Δ*L*_i_ point to structural elements that are important for inducing communication. Looking at the respective Δ*L*_i_ profiles in more detail, it is interesting to note a differential degree of change in some regions depending on the ligand bound configuration of the NTD. Residues residing in ß-strand_1_ (res 15-26) and the ATP-lid (res 112-136), residues D302, S330-L335, helix_10_ (res 407-427), and residues neighbouring Y604 and Y627 all record sharp positive Δ*L*_i_ in the presence of ADP, and negative changes for ATP. ß-strand_1_ swapping between protomers contributes to the stabilization of NTD dimerization (Ali *et al*, 2006; Dollins *et al*, 2005; Pearl & Prodromou, 2006), and its deletion has been shown to affect ATPase activity in mitochondrial and yeast Hsp90 homologs (Richter *et al*, 2002; Lavery *et al*, 2014). The ATP-lid is intricately implicated in the progression of conformational change, acting as a trigger for NTD dimerization in response to bound ATP (Ali *et al*, 2006; Pearl & Prodromou, 2006; Schulze *et al*, 2016; Prodromou *et al*, 2000). Helix_10_ has been previously implicated in nucleotide driven allosteric signal propagation between the NTD and M-domain in *E. coli* HtpG (Seifert *et al*, 2012). Residues Y604 and D621-Y627 reside in helix_18_ and helix_19_ respectively, and undergo large negative Δ*L*_i_ in the ATP and apo complexes, but display positive Δ*L*_i_ in the ADP complex, suggesting a tighter binding of these interface elements in the ATP and apo complexes. This observation is in agreement with the conformational dynamics analysis of the PO structures in which interactions between these residues from either protomer are lost in the presence of ADP, but are retained in the ATP and apo complexes.

Studying the respective Δ*L*_i_ profiles for the FO complexes (**Figure 7**) reveals largely negative Δ*L*_i_ for FO-ATP, while Δ*L*_i_ for FO-ADP and FO-apo fluctuate. Notable differences are observed between the protomers of each complex. Large contrasting Δ*L*_i_ values are recorded in the NTD ß-stand_1_, helix_1_, and the ATP-lid regions, in which protomer 1 records positive change and protomer 2 a negative change. This differential behaviour may be further evidence of previously reported asymmetrical protomer dynamics (Flynn *et al*, 2015; Mayer & Le Breton, 2015). As seen in the closed complexes, certain regions of the protein respond differentially in response to the bound configuration of the NTD. Here, residues E200-K224, forming ß-strand_8_ located at the interface between the NTD and the charged linker, experience negative Δ*L*_i_ in the ATP complex, and positive Δ*L*_i_ for the ADP and apo configurations, demonstrating a reduced flexibility in this region when ATP is present. In fact, these residues have been directly implicated in the regulation of the human Hsp90α ATPase cycle, whereby mutations to select residues in this region lead to increased NTD flexibility, resulting in severely altered chaperone activity and a shift in the conformational equilibrium in favour of the open state (Tsutsumi *et al*, 2009). In the M-domain, residues R367-E380 and L396-Q404 are in close spatial proximity to one another, and all experience sharp positive Δ*L*_i_ in the presence of ADP compared to ATP. Positioned at the NTD / M-domain interface, they have also been reported to form a nucleotide sensitive hinge, implicated in allosteric signal propagation from the NTD to the M-domain (Morra *et al*, 2012; Blacklock *et al*, 2014a). In the CTD, both helix_18_ and helix_19_ experience positive Δ*L*_i_ in all three complexes; however, FO-ADP experiences greater change, once again suggesting looser interactions under these conditions.

**Figure 7:**
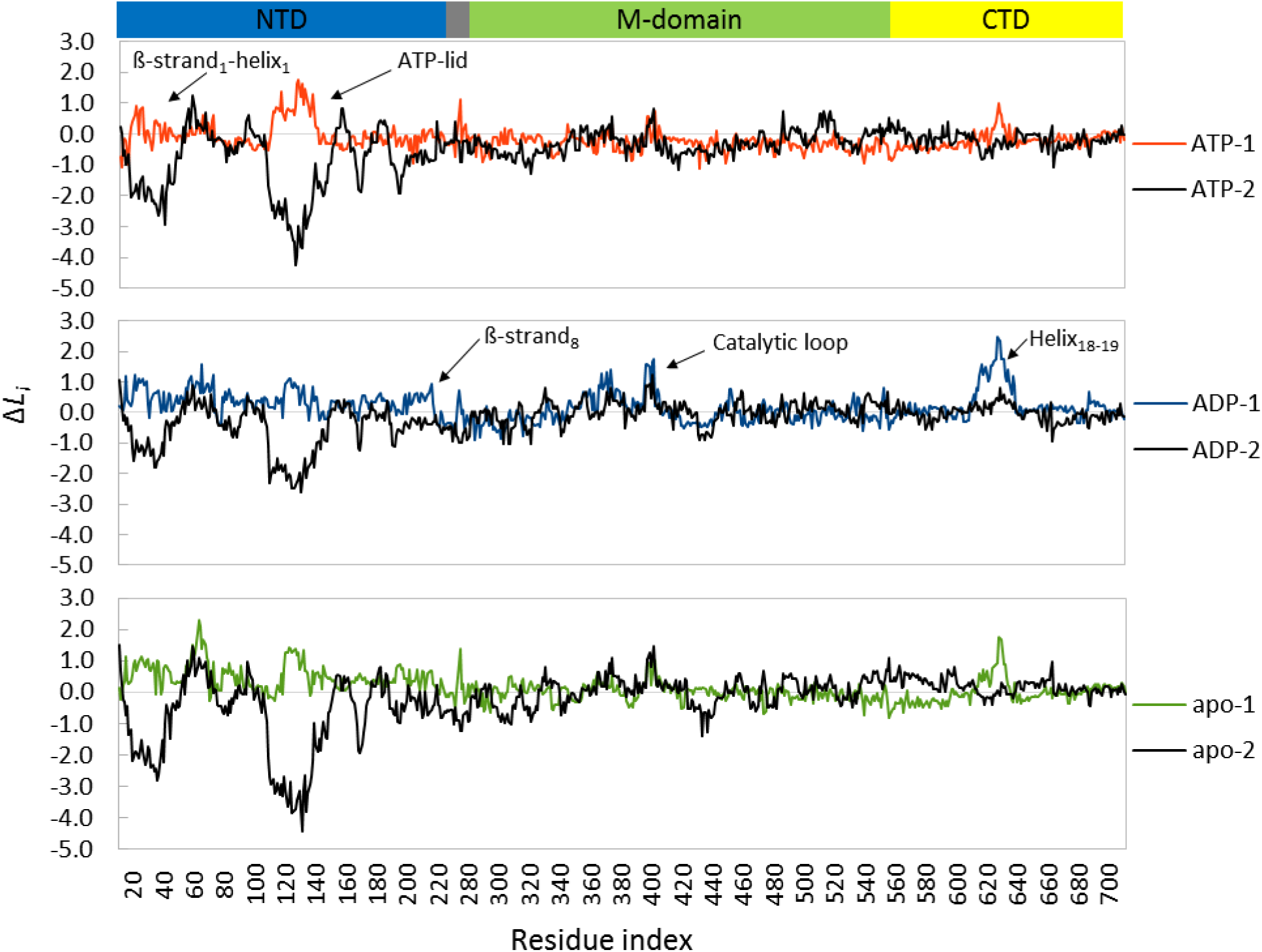
Change in reachability (*ΔLi*) of each residue for the fully-open complexes. Protomer 1 is coloured by NTD configuration (ATP – red, ADP – blue, apo – green), and protomer 2 is in black.

Overall, Δ*L*_i_ provides a unique metric for monitoring the effect of nucleotide modulation on local and global restructuring and its implication on long range communication in Hsp90, highlighting regions of the protein that may be involved in steering conformational change. Here, the analysis confirms the tensing effect of ATP, with the protomers of both conformations experiencing negative Δ*L*_i_ values. Conversely, protomer relaxation afforded by ADP binding is also confirmed, with the average path length increasing in both ADP bound conformations. Regions that experience large Δ*L*_i_ are most notable in the open complexes, and include residues belonging to the functionally important ß-strand_1_, helix_1_, ATP-lid, and catalytic loop (Ali *et al*, 2006; Pearl & Prodromou, 2006; Prodromou *et al*, 2000), as well as the helix_18_/helix_19_ CTD interface.

#### Betweenness centrality points to putative communication hubs

BC has been shown to be an effective measure for identifying functional residues implicated in the control of intra-protein communication (Ozbaykal *et al*, 2015; Brown *et al*, 2017b), as well as protein-ligand (Liu *et al*, 2011) and protein-protein binding sites (del Sol *et al*, 2005; Cheng *et al*, 2007) (see review (Taylor, 2013) for other examples). Here, BC is calculated for the closed and FO complexes to identify individual/groups of residues important for intra- and inter-protomer communication.

The respective BC profiles of both conformations (**Figure 8**), reveal that bound nucleotide has little impact on modulating communication hubs in either conformation, with similar trends observed for all three configurations. Thus, to simplify data presentation, the average BC over all three configurations for each conformation is shown (**Figure 8** black curves). Peak locations in each conformation describe residues with high BC and thus high frequencies of usage in intra-protein communication. As previously mentioned, BC for a given residue *i* is a measure of the number of shortest paths that visit *i*. In the closed complexes, the number of shortest paths is expected to exceed that of the FO complexes due to compactness of structure, leading to increased intra-and inter-protomer communication and thus a greater number of communication hubs. The peak residues of the closed complexes are tightly grouped, listing residue regions rather than select residues as seen in the open complexes (**Figure 8**-A & B). Nevertheless, both conformations share several overlapping peak residues that correlate well with known functional residues, particularly those involved in regulating ATPase activity and conformational dynamics. Peak residues listed by BC are summarized in

**Figure 8:**
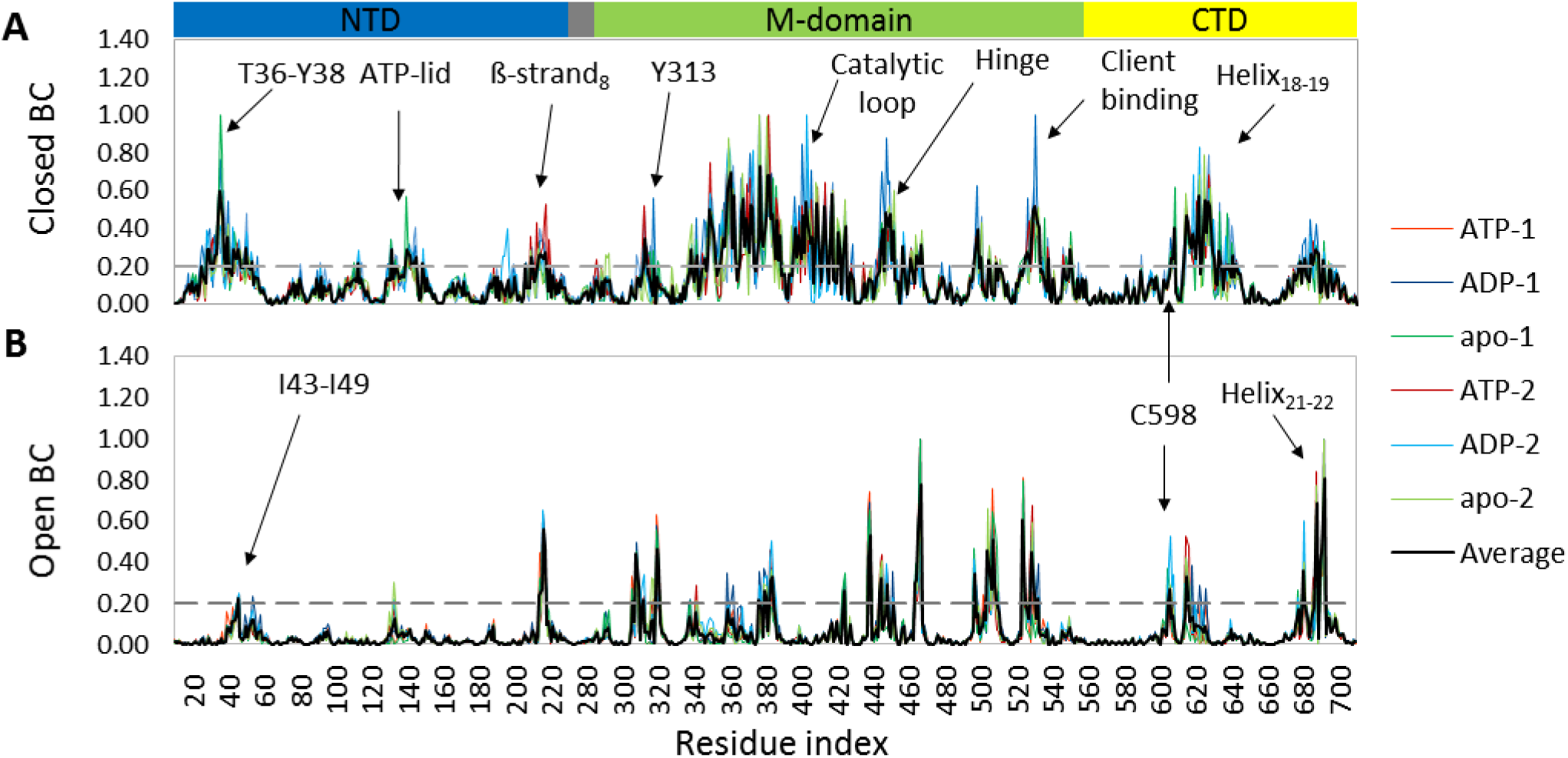
Betweenness centrality (BC) profiles: Showing regions in the (A) closed and (B) open complexes that may constitute putative communication hubs. Dashed grey lines indicates the peak residue threshold.

**Table 2** and mapped to their respective 3D structures in **Figure 9**. Their respective functional importance is discussed below.

**Table 2:**
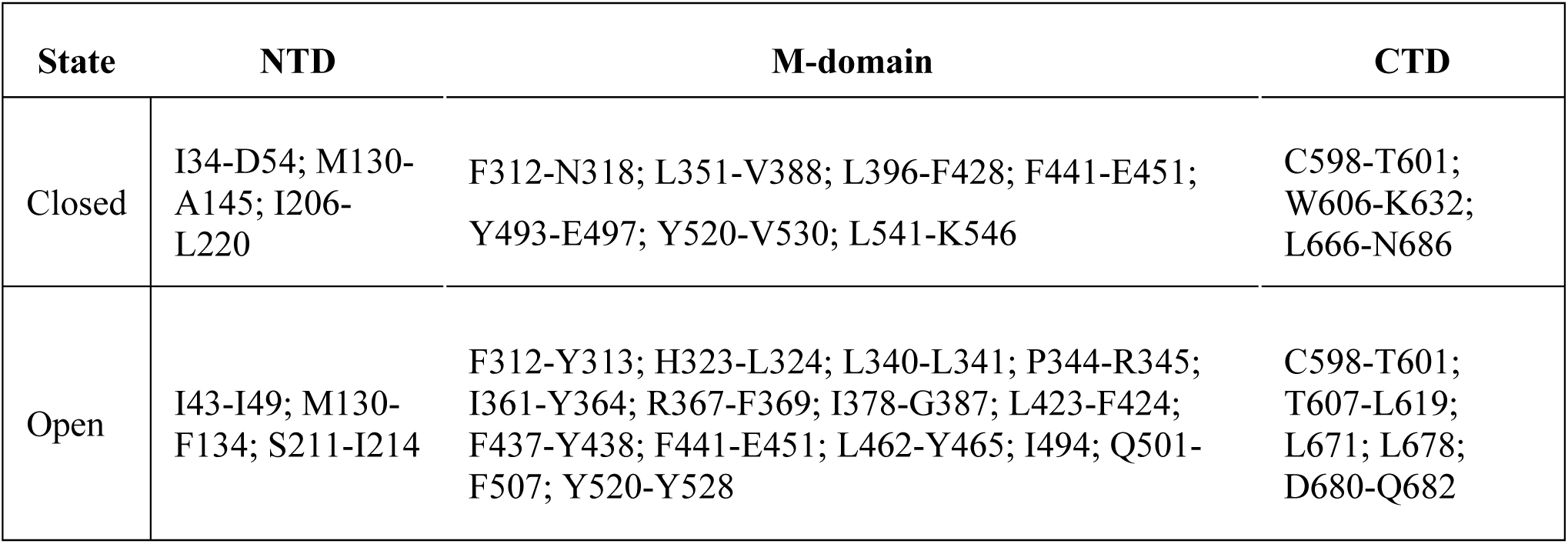
Summary of residues with high BC for the closed and open states

| State | NTD | M-domain | CTD |
| --- | --- | --- | --- |
| Closed | I34-D54; M130-A145; I206-L220 | F312-N318; L351-V388; L396-F428; F441-E451; Y493-E497; Y520-V530; L541-K546 | C598-T601; W606-K632; L666-N686 |
| Open | I43-I49; M130-F134; S211-I214 | F312-Y313; H323-L324; L340-L341; P344-R345; I361-Y364; R367-F369; I378-G387; L423-F424; F437-Y438; F441-E451; L462-Y465; I494; Q501-F507; Y520-Y528 | C598-T601; T607-L619; L671; L678; D680-Q682 |

**Figure 8:**
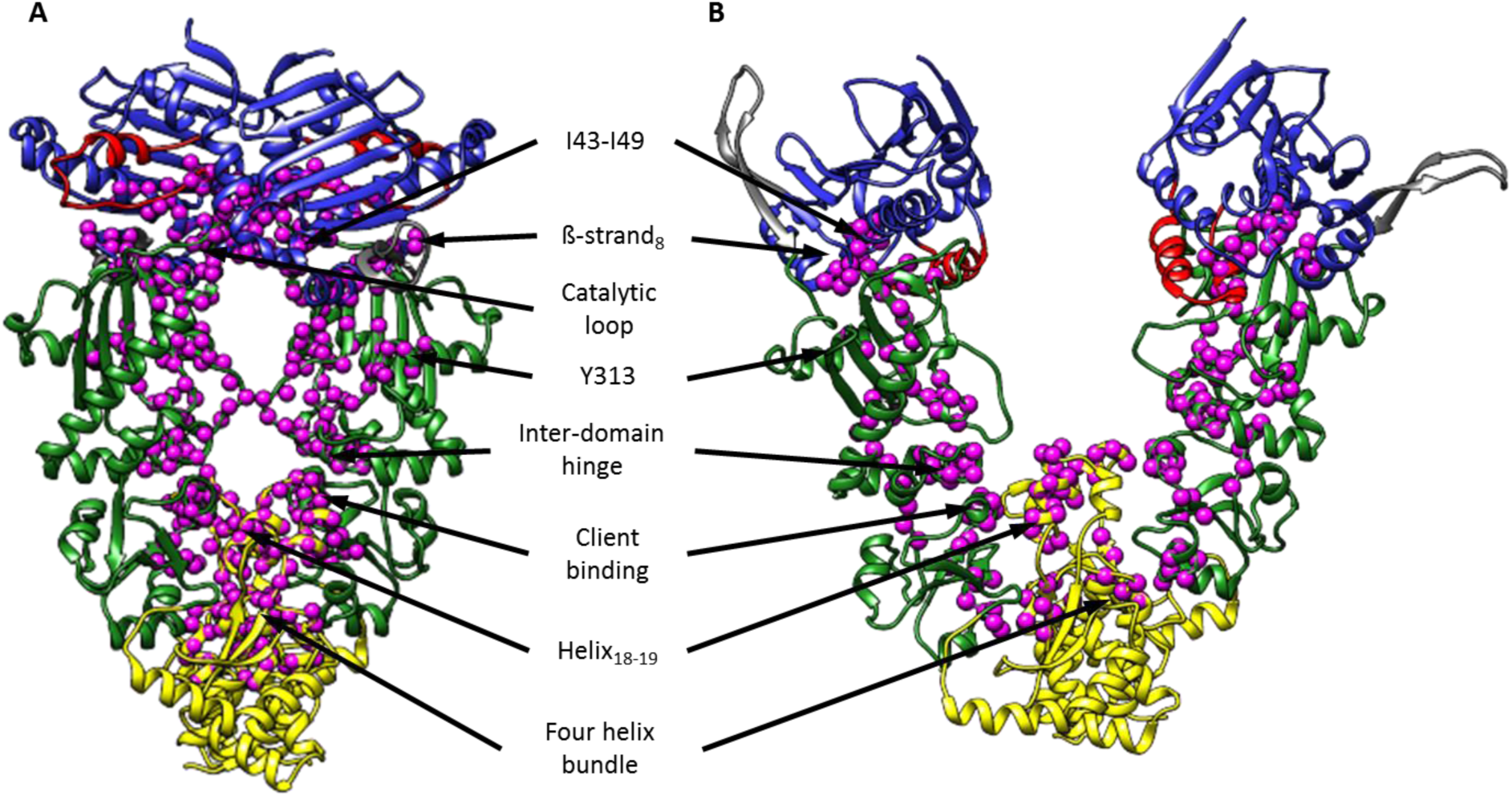
Structural mapping of high BC residues. (A) Closed complex and (B) fully open complex. Showing peak residues as magenta spheres.

Starting with the NTD, phosphorylation and mutation studies have linked T36 and Y38 to ATPase regulation and altered binding of co-chaperones Aha1 and Cdc37 (Mollapour *et al*, 2011; Prodromou *et al*, 2000; Cunningham *et al*, 2008), while computational studies have implicated both residues in inter-domain signal propagation (Neckers, 2002; Retzlaff *et al*, 2009; Morra *et al*, 2009). These residues are only selected in the closed complexes where they help stabilize NTD dimerization by contributing to crucial interactions with the catalytic loop (S391-S406), and residues R46-I49 (Ali *et al*, 2006; Meyer *et al*, 2003). Residues I43-I49 are listed in both conformations, and mutation studies have shown that E47A abolishes ATP hydrolysis suggesting a catalytic function *in vivo* (Obermann *et al*, 1998). In the closed conformation, E47A would have a negative impact on catalytic loop interactions and thus decrease ATPase activity. In the open state, I43-I49 are positioned in closed proximity to the open ATP-lid, presumably helping to stabilize it in the configuration. Furthermore, BC also lists lid residues M130-F134 in both conformations. In the open complex, these residues are uniquely positioned over I43-I49, while in the closed complex, they are positioned over residues S53 and D54, which together form a hydrophobic patch known to be involved in ATP-lid stabilization (Prodromou *et al*, 2000). In sum, these findings point to I43-I49 and M130-F134 as crucial communication hubs in both conformational states.

Next, residues I206-L220 of ß-strand_8_ and helix_7_ are located at the interface between the NTD and charged linker and have been implicated in chaperone secretion and modulation of the ATPase where I218A and L220A mutations are thought to alter the charged linker’s topology, causing reductions in co-chaperone association and *in vivo* cell death (Tsutsumi *et al*, 2009). Of these residues, only S211-I214 are listed in both conformations, and we note that in the closed complexes, ATP records higher BC values for these residues than the ADP and apo configurations. The apparent loss of BC in the ADP and apo complexes is significant in that the mutation I218A has also been shown to impact conformational dynamics, shifting the conformational equilibrium in favour of the open state (Hessling *et al*, 2009). It is possible that I206-L220 may act as a crucial communication hub capable of sensing bound nucleotide at the NTD. Under rigid ATP conditions I206-I220 may help stabilize the complex in a closed conformation. Eventual ATP-hydrolysis and subsequent protomer relaxation in response to ADP binding may destabilize this region, which triggers a conformational shift to the open sate.

Looking at the listed residues in the M-domains of both conformations, phosphorylation of Y313 has been shown to promote Aha1 binding affinity by inducing structural rearrangements that favour co-chaperone binding (Xu *et al*, 2012). Residues V368-I370 have been previously identified as key interdomain contacts in a comprehensive network analysis of yeast Hsp90 (V384-I350) (Blacklock *et al*, 2014a). Of these residues, the F369A mutation in yeast has been found to drastically decrease ATPase activity by disrupting the positioning of the catalytic R400 which in turn leads to destabilization of the ATP-lid (Siligardi *et al*, 2004; Schulze *et al*, 2016; Meyer *et al*, 2003). Residues L396-L409, listed in the closed but not the open complexes, represent the M-domain catalytic loop where they are thought to form an important interdomain hinge between the NTD and M-domain (Morra *et al*, 2012; Rehn *et al*, 2016). In addition to the this, S399 (Soroka *et al*, 2012), R400 (Meyer *et al*, 2003), E401 (Panaretou *et al*, 2002), and Q404 (Meyer *et al*, 2003) have been implicated in modulating ATPase activity and signal propagation (Morra *et al*, 2009; Blacklock *et al*, 2014a). The aromatic cluster formed by F384 and F441 are listed by BC in both conformations, and are thought to be important as allosteric control points for nucleotide dependent conformational change, acting as a mechanical hinge between the M-and CTDs (Morra *et al*, 2012; Rehn *et al*, 2016). Reside E451 is also selected in both conformations and mutation studies have shown that E451A decreases inter-protomer interactions leading to decreased ATPase activity (Rehn *et al*, 2016), while E451K is thought to perturb the M-domain structure impacting gluticoid receptor binding (Street *et al*, 2014). Phosphorylation mimicking mutations to S505 is fatal for yeast viability (S485) and is thought to impart resistance to ATP induced conformational transition to the closed state, favoring the open state (Soroka *et al*, 2012). Here, S505 is only selected in the open complexes suggesting an important role in this conformation, possibly as a post translation modification site for regulating the closing transition. Residues E527, Y528, and Q531 have been shown to form part of an important region for client binding in both yeast and *E. coli* Hsp90s (Genest *et al*, 2013) and have also been implicated in allosteric regulation (Blacklock *et al*, 2014b), while T545I mutation has been shown to decrease client binding in *E. coli* (Street *et al*, 2014).

In the CTD, BC lists several prominent peaks in both conformations: The C598-T601 peak includes C598 which has been heavily implicated as a crucial allosteric switch, capable of regulating ATPase cycle (Retzlaff *et al*, 2009; Morra *et al*, 2009). Peak T607-L619 in the open conformation corresponds to helix_18_ at the CTD interface. In the closed conformation this peak is extended to include residues W606-K632 representing a coiled loop helix18-19. Of these residues: W606 is thought to be involved in client binding and chaperone activity (Genest *et al*, 2013); phosphorylation mimicking mutations to S623 and M625 affect inter-subunit communication leading to decreased ATPase rates in yeast (Soroka *et al*, 2012); while phosphorylation of Y627 is thought to induce structural rearrangements that reduce the binding affinity of Aha1 (Xu *et al*, 2012). Finally, several residues at the four helix bundle (helix21-22) located at the extreme C-terminal dimerization site are selected in both conformations. Included in this peak is D680 which has been shown to play an important role in maintaining inter-protomer contacts (Rehn *et al*, 2016), while several residues from this site in yeast Hsp90 have been reported to participate in fast communication with residues in the NTD (Morra *et al*, 2009), suggesting a nucleotide sensitive regulatory role.

Comparatively, BC selects several groups of residues in both the closed and open structures regardless of bound nucleotide, suggesting key intra-and inter-protomer communication points. These regions correspond to known functional sites of which several are implicated in conformational dynamics. Furthermore, BC also list several residues in regions of unknown function.

### Perturbation response scanning

PRS is a technique used to assess the allosteric influence each residue has on all other residues when externally perturbed. Several previous studies have demonstrated the use of PRS as an effective approach for the identification of functionally important sites that are potentially involved in modulating binding-region motions (Atilgan & Atilgan, 2009; Atilgan *et al*, 2010; General *et al*, 2014; Abdizadeh *et al*, 2015; Abdizadeh & Atilgan, 2016; Penkler *et al*, 2017), and PRS analysis has been previously shown to be an efficient method for determining the allosteric potential of large multi-domain proteins (General *et al*, 2014; Penkler *et al*, 2017). Briefly, PRS uses linear response theory to predict whole protein displacements in response to the insertion of external forces at single residue sites (see equation (2). In this study, 250 perturbations were sequentially applied to each residue in the protein and the resultant displacement of the whole protein in response to each perturbation recorded in a 3D matrix and analysed in terms of: (1) average relative displacement-to gain insight on protein sensitivity to external force perturbation, and (2) conformational overlap with a known target state-to assess the potential of each residue to select displacements that are representative of an expected conformational change. Together these analyses provide insight regarding site on the protein that are potentially involved in modulating conformational dynamics.

#### Average residue displacements identify putative allosteric effector and sensor residues

Analysis of the average response of the whole protein to 250 perturbations of each residue allows for the identification of sites on the protein that are allosterically sensitive to external force perturbations. Conversely, residues whose perturbations result in large conformational changes elsewhere in the protein can be seen as allosteric effectors, representing sites on the protein likely involved in external binding events such as specific co-chaperone and client binding in the case of Hsp90. This analysis approach for PRS has been previously demonstrated on Hsp70 by General *et al,* (2014). The average displacement response maps for both configurations of the protein are shown in **Figure 10**, in which the *ik*^th^ entry refers to the average response (displacement) of residue *k* to external perturbation at residue *i*. Thus, bright spots indicate peak residues that experience large displacements, while darker areas imply residues that experience little or no displacement. Accompanying each map, are the most influential ‘effector’ residues (rows) in the right hand bar plots, and the most sensitive ‘sensor’ residues (columns) in the lower column plots,

For the closed conformation complexes (**Figure 10**-A), comparing the displacement intensities of the different NTD configurations reveals the ADP complex to be the most sensitive to external force perturbations, while the apo complex appears to be the least sensitive. This observation may suggest that the ADP closed complex is more amenable to conformational change compared to the other configurations, in agreement with the hypothesis that ADP imparts protomer flexibility. Strong sensor and effector signals are recorded at the linker regions of all three complexes. These residues correspond to the Gly-Gly-Gly-Gly insert, introduced in the homology modeling process to cover the missing linker region (see methodology), and their high degree of displacement is to be expected as glycine residues are known to be highly mobile elements. Several sensor peaks are observed in all three closed configuration complexes: ß-strand_1_ (res 15-20); helix_1_ (res 21-36); the first turn of the NTD ß-sheet (res 81-88); the ATP-lid (res 120-129); ß-strand_4_ (res 170-180); the loop connecting helix_12_ and helix_13_ (res 469-472); as well as helix_17_ (res 550-590) and helix_20_ (res 638-660) in the CTD. Nucleotide dependent differences are also observed for residues that correspond to Aha1 binding (res 300-308) (Meyer *et al*, 2004), the catalytic loop and interdomain hinge (Morra *et al*, 2012) (res 391-406), the phosphorylation site Y313 (Xu *et al*, 2012), helix_13_ (res 478-490), and the putative client binding site at helix_16_ (res 523-530) experience increased sensitivity in the presence of ADP, but not the ATP and apo complexes. Studying the corresponding effector sites, inter-protomer coupling is observed between the NTD and CTD in all three complexes, where perturbations at the CTD (helix_17_ and helix_20_) of protomer 2 are strongly detected at the NTD of protomer 1 and *vice versa* (**Figure 10**-A, white boxes). For the ATP complex, NTD perturbations in protomer 1 appear to illicit a stronger CTD response in both protomers compared to the ADP and apo complexes, while CTD perturbations in protomer 2 of the ADP and apo complexes appear to illicit a stronger response at the NTD of protomer 1 compared to the ATP complex. This observed allosteric coupling between the two terminals is in agreement with a previous computational study demonstrating efficient allosteric communication between the NTD and CTD in response to bound nucleotide (Morra *et al*, 2009). Other notable effector residues include the ATP-lid (res 121-126), particularly in the ADP bound complex, helix_10_ (res 406-425), and residues surrounding A469 and G515 in the M-domain. These sites are significant in that ATP-lid stabilization is necessary to maintain the closed ‘active’ state, while helix_10_ has been implicated in allosteric signal propagation between the NTD and M-domain in HtpG (Seifert *et al*, 2012). Furthermore, all of the M-domain sites are located on the surface of the protein, in positions easily accessible to co-factor binding.

**Figure 10:**
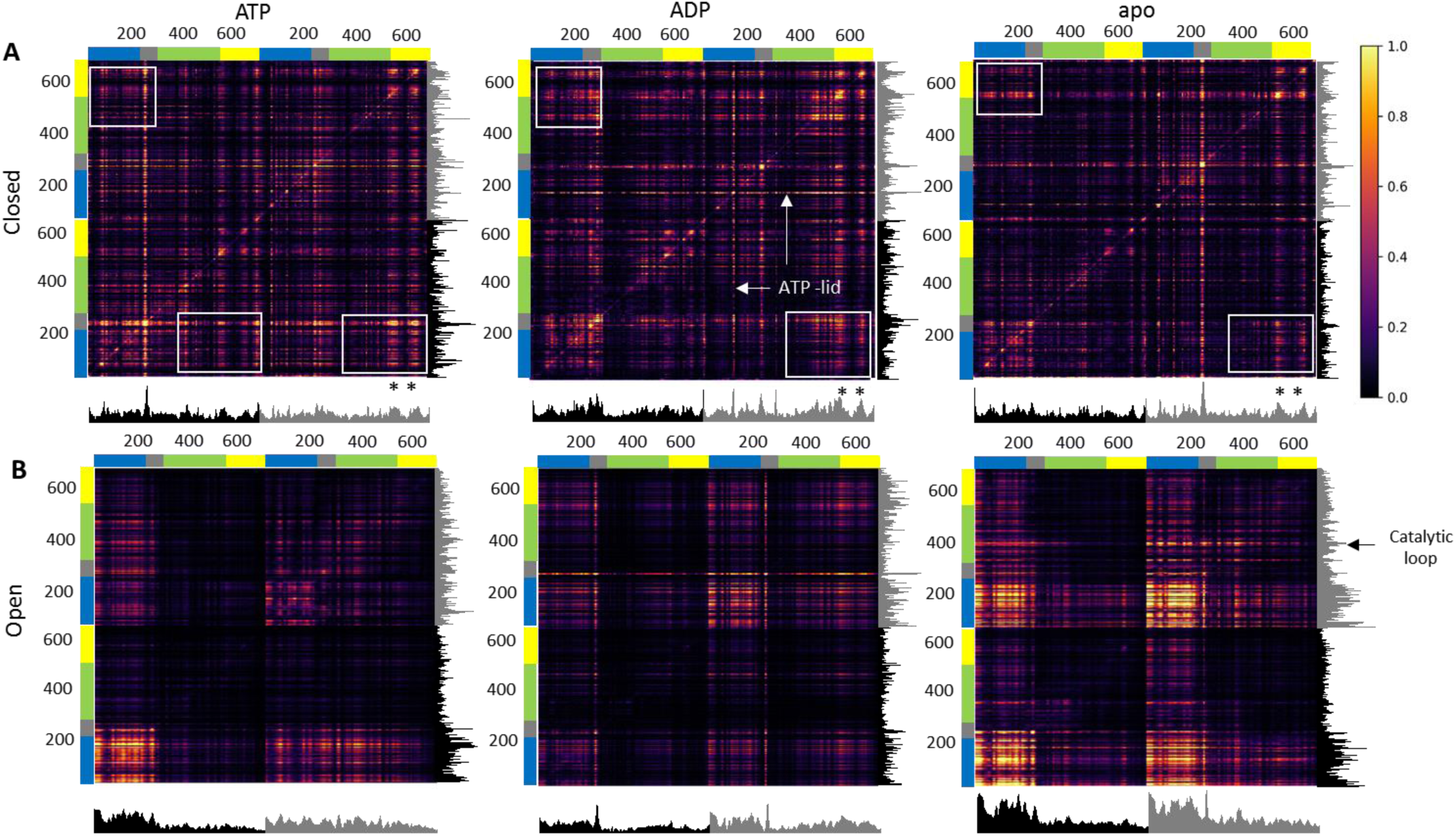
PRS response maps showing effector and sensor regions of the protein. (A) Closed and (B) fully open complexes. Columns denote sensor residues, while rows represent effector (perturbed) residues. Bright spots indicate highly sensitive residues that experience large displacements. Accompanying each map are ‘effector’ bar plots on the right-hand-side and at the bottom ‘sensor’ column plots (Protomer 1-black, Protomer 2-grey). (*) Denotes helix_17_ and helix_20._.

The open conformation complexes (**Figure 10**-B) show clear differentiation between the different complex configurations. However, all three complexes show strong allosteric coupling between their NTDs, where perturbations at the NTD of one protomer lead to strong displacement signals at NTD of the second protomer. Interestingly, the apo complex appears to be the most sensitive to NTD perturbations while the ATP complex appears to be more sensitive than ADP. This observation suggests the apo complex to be more amenable to conformational change, an observation that is in agreement with previous reports describing stochastic conformational dynamics for apo Hsp90 (Krukenberg *et al*, 2008, 2011; Southworth *et al*, 2008). Looking in more detail, it is also apparent that perturbations arriving at the NTD result in displacements at the M-and CTD depending on the nucleotide configuration. M-domain displacements at ß-turn E332-E336 and the catalytic loop (res 395-407) are observed within the perturbed protomer of the ATP and apo complexes, while CTD sensitivity at helix_17_ (552-570) is observed for the ADP complexes. Interestingly, most of these nucleotide specific sensor sites are also listed as allosteric effectors for the respective complexes, implying allosteric coupling between these sites.

#### Perturbation of key residues reveals sites likely involved in modulating conformational change

In this section, PRS is used to identify residues whose perturbation invoke a conformational change towards an expected target conformation representative of the opposite state. For each residue, the displacement response for each sequential perturbation is compared to a known experimental displacement (opening or closing), and the goodness of fit measured using the Pearson’s correlation coefficient (equation (3). In this manner, each residue records 250 correlation coefficients, and the highest correlation, *C*_*i*_, is selected to represent the maximum potential for that residue to invoke a conformational change towards the opposite state. The closed complexes are assessed in terms of the opening transition, and the FO complexes in terms of the closing transition. The results of this analysis are shown in **Figure 11** and **Figure 12** for the closed and FO complexes respectively, and the highest *C*_*i*_ residues are summarized in **SD 8**.

**Figure 11:**
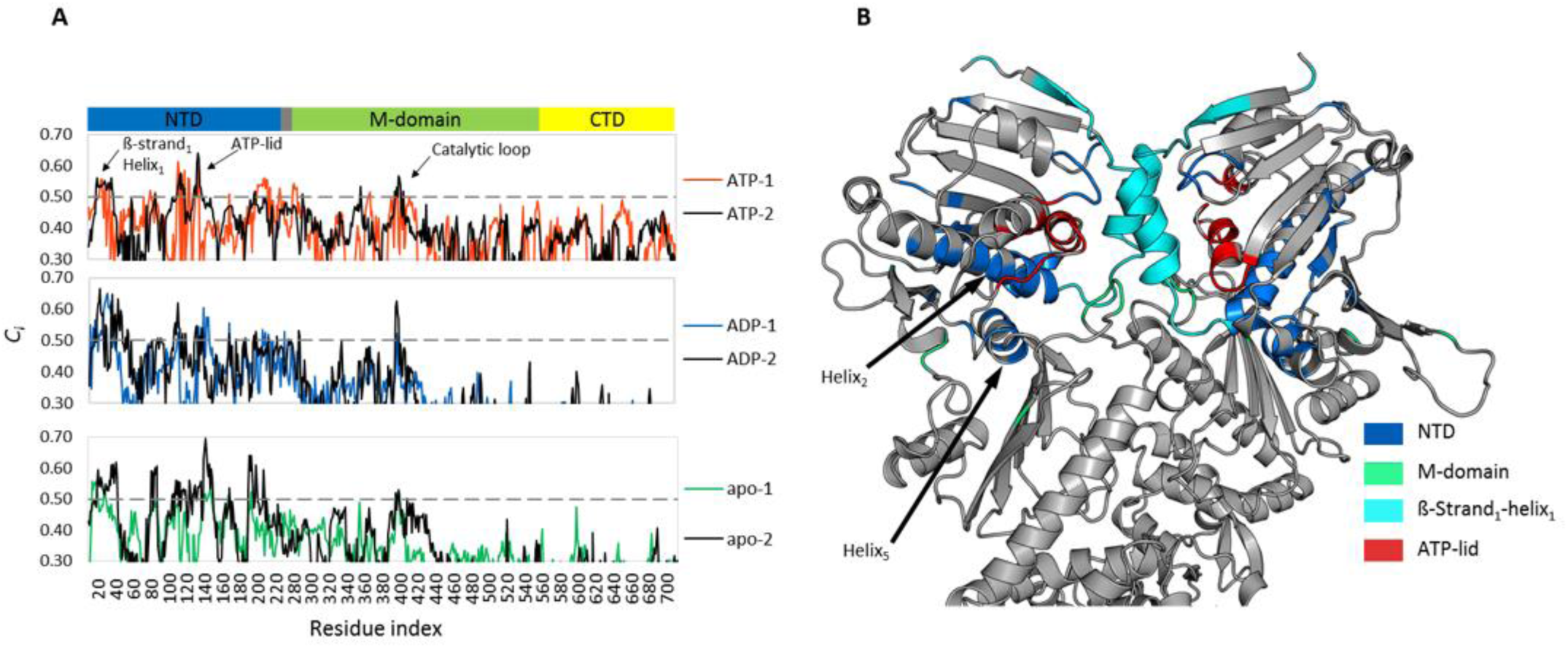
Residues accentuated by PRS in the closed conformation complexes. (A) PRS profiles showing accentuated residues which select the open conformation. Protomer 1 is coloured by NTD configuration (ATP–red, ADP–blue, apo–green), and protomer 2 in black. (B) Zoomed in view of the NTD of the closed conformation, showing the accentuation of the NTD dimerization interface of the ADP bound complex. Residues accentuated by PRS are colored according to domain location.

**Figure 12:**
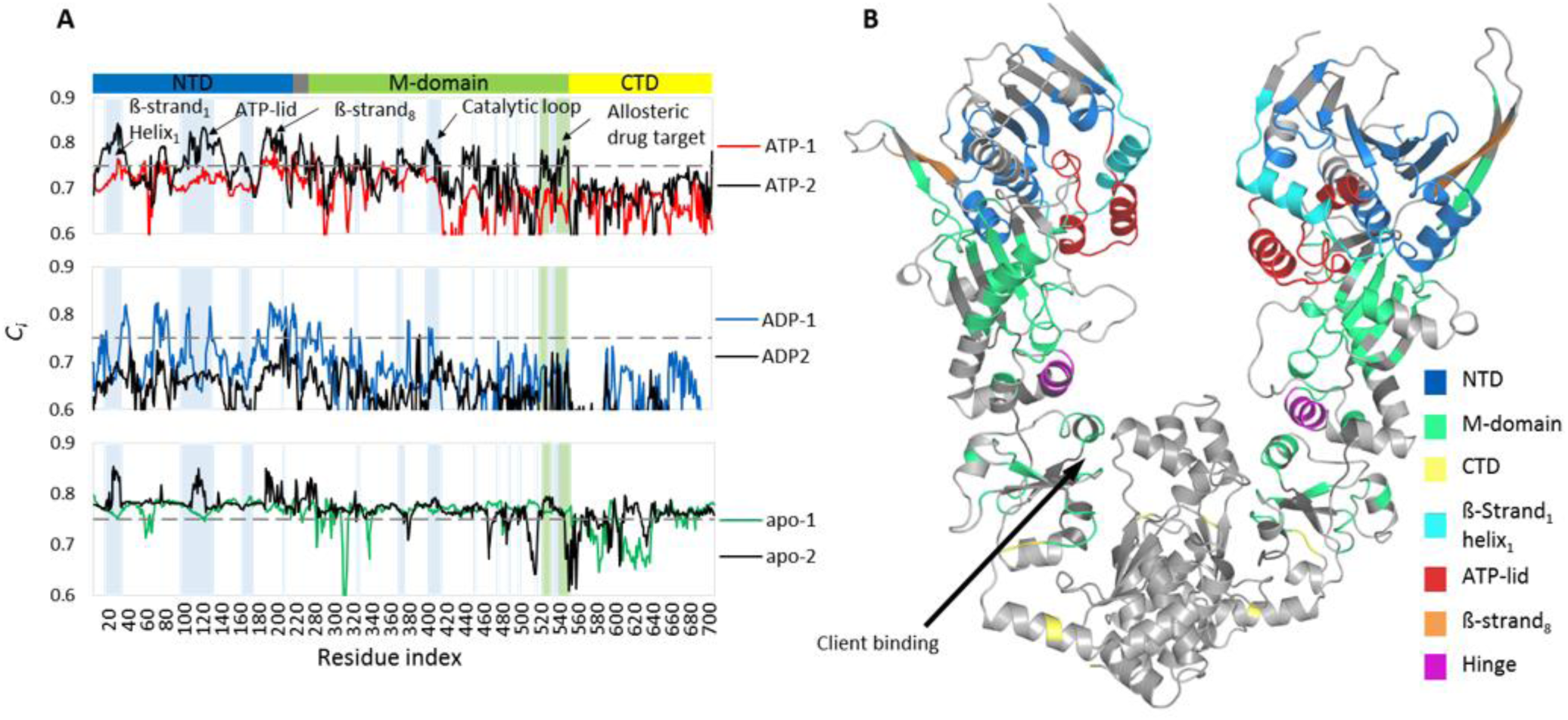
Residues accentuated by PRS in the open conformation complexes. (A) PRS profiles showing accentuated residues capable of selecting the closed conformation. Protomer 1 is coloured by NTD configuration (ATP–red, ADP–blue, apo–green), and protomer 2 in black. Blue shading represents overlap with Aha1 binding residues and the green shading client binding residues. (B) Structural mapping of residues accentuated by PRS for the FO-ATP complex coloured by location.

For the closed conformation complexes (**Figure 11**-A), PRS indicates that regardless of bound nucleotide, no single residue is capable of selecting the open conformation in response to external force perturbation, with peak residues recording maximum *C*_*i*_ values of ∼0.60. Despite this, several *C*_*i*_ residue peaks (*C*_*i*_ > 0.50) appear to correspond to functional sites previously identified and discussed in the BC analysis. Accentuated residues belonging to ß-strand_1_ (res 16, 19, 20); helix_1_ (res 23-40); the ATP-lid (res 107-113, 116, 119, 130-132, 134, 135, 137-139, 142-145); and the catalytic loop (res 397-400) form a central dimerization hub at the NTD and have been previously described as forming part of an NTD/M-domain mechanical hinge (Morra *et al*, 2012) (**Figure 11-**B). Destabilization of these inter-protomer interfaces is crucial before protomer uncoupling and a conformational shift to the open state can occur (Cunningham *et al*, 2008). PRS also accentuates residues residing in helix_2_ (res 41-65); ß-strand_2_ (res 88-94); helix_5_ (res 203, 205, 209-211); and ß-strand_8_ (214, 219). Of these, helix_2_ has been previously reported to play a crucial role in propagating allosteric signals from the NTD to the M-domain in *E. coli* Hsp90 (Seifert *et al*, 2012), while residues residing in ß-stand_2_ have been associated with long range communication with the CTD (Morra *et al*, 2009). The NTD / linker interface formed by helix_5_ and ß-strand_8_ has been implicated in modulating chaperone affinity (Tsutsumi *et al*, 2009) and conformational transitions in response to bound nucleotide (Hessling *et al*, 2009). Comparing the highest *C*_*i*_ between complexes, residues belonging to the ATP-lid recorded the highest *C*_*i*_ values in the ATP and apo complexes (0.62 and 0.67 respectively), while residues belonging to ß-strand_1_, helix_1_, and the catalytic loop recorded the highest *C*_*i*_ values for the ADP complex.

Studying the open complexes, the nucleotide configuration appears to have a marked differential effect on the potential of each complex to interconvert to the closed state. Comparing the percentage of residues with *C*_*i*_ > 0.75, the ATP (33%) and ADP (11%) complexes record far fewer than the apo complex (85%). The elevated number of *C*_*i*_ > 0.75 in the apo complex is interpreted to be indicative of a natural propensity for this complex to interconvert to the closed state, an observation in keeping with the hypothesis that the apo protein behaves stochastically (Krukenberg *et al*, 2008, 2011; Southworth *et al*, 2008). In the case of the nucleotide bound complexes, PRS only lists select residues capable of eliciting a conformational transition towards the closed state.

Looking at the *C*_*i*_ profile for the ATP bound complex (**Figure 12**-A), PRS accentuates several residue clusters in the NTD and M-domain (**Figure 12**-A, peaks) which once again point to functionally important regions. In the NTD, these residues include: ß-strand_1_ (res 20-24); helix_1_ (res 25-35); helix_2_ (res 43-53); the ATP-lid (res 113-136); ß-strand_5_ (res 163-170, 173-174); ß-strand_6_ (res 186-190); helix_6-7_ (res 191-214); and ß-strand_8_ (res 219-224), and in the M-domain: helix_9_ (res 312-316); several residues belonging to the ß-sheet (res 319, 320, 326, 330-338, 341, 343, 347, 360, 361, 363, 376-393); the catalytic loop (res 402-410); helix_10_ (res 411, 412, 416-418, 420); helix_12_ (res 443-449, 451-454); helix_16_ (res 523-525, 527, 529, 530); and lastly loop residues T540-L551. Of these accentuated sites, the functional importance of ß-strand_1_, helix_1-2_, ß-strand_8_, the ATP-lid, and the catalytic loop have already been discussed in context of the opening transition; however, accentuation of these elements by PRS for the FO-ATP complex, and thus the closing transition, also bears functional significance. It is widely believed that ATP binding initiates the closing transition by triggering the uncoupling of the ß-strand_1_ and helix_1_ from the ATP-lid, leading to its release and closure over the nucleotide binding pocket. This conformational repositioning of the ATP-lid not only entraps bound nucleotide, but also facilitates N-terminal dimerization through exposure of a hydrophobic surface required for ß-strand_1_ swapping and stabilization of the closed active complex (Cunningham *et al*, 2008). In addition to these NTD conformational dynamics, ATPase activation of the closed complex is only fully achieved when the M-domains dock onto their respective NTDs, allowing the M-domain catalytic loop to be repositioned such that R400 projects into the nucleotide binding pocket to coordinate with the γ-phosphate of ATP (Meyer *et al*, 2003; Prodromou, 2012). Interestingly, we note that PRS accentuates ß-strand_1_, helix_1_, and the ATP-lid in the FO-ATP complex, but not in the FO-ADP complex (**Figure 12-**A), a finding that supports the hypothesis that ATP binding allosterically activates the closing transition, while bound ADP stabilizes the open conformation. The conformational dynamics required for ATPase activation in Hsp90 are thought to be the rate limiting step in an inherently slow process, recording rate constants in the order of minutes (Prodromou *et al*, 1997; Scheibel *et al*, 1997; McLaughlin *et al*, 2002; Panaretou, 1998). Indeed, a recent study has linked the rate constant of ATP hydrolysis with the rate constants of ATP-lid closure, ß-strand swapping, and intra-protomer association of the NTD and M-domains (Schulze *et al*, 2016). Furthermore, the authors of this study propose a two-step mechanism for ATP-lid closure and find evidence of a previously unknown mode of action for the co-chaperone Aha1, where they suggest its involvement in facilitating early mobilization of the ATP-lid (Schulze *et al*, 2016). Here, we find that residues accentuated by PRS in the ATP complex are clustered around NTD and M-domain sites that have been previously implicated in Aha1 binding (Meyer *et al*, 2003; Retzlaff *et al*, 2010) (**Figure 12**-A, blue shading). We propose that perturbations arriving at ß-strand_1_, helix_1_, and the ATP-lid may occur naturally through binding interactions with Aha1, providing possible evidence in support of Aha1 assisted ATP-lid closure and interstate conversion to the closed state. Also of functional importance and accentuated by PRS in the ATP complex, are residues belonging to helix_12_ corresponding to the M-domain/CTD hinge (Morra *et al*, 2012; Blacklock *et al*, 2014a), as well as helix_16_ and loop T540-L551 which are thought to be important regulatory sites for client binding (Genest *et al*, 2013; Street *et al*, 2014; Blacklock *et al*, 2014a) (**Figure 12**-A, green shading). Interestingly, this CTD region has been previously reported to be an allosteric drug target site (Morra *et al*, 2010), and subsequent drug discovery studies have demonstrated how bound ligands at this site can allosterically enhance ATPase activity through the asymmetric modulation of protomer conformational dynamics (Sattin *et al*, 2015; Vettoretti *et al*, 2016).

Overall, the data presented here suggests PRS to be a suitable technique for identifying co-factor binding sites that may be specifically involved in allosteric modulation of conformational dynamics. To tests this hypothesis, we analysed the PRS data for the FO-ATP complex in terms of sites known to be involved in binding interactions with the heat shock organising protein (HOP) (res: E307, E311, K314, N318, W320, D322, K431, E432, E473, A469, T482, and E486) (Schmid *et al*, 2012; Hatherley *et al*, 2013). HOP is thought to modulate Hsp90’s conformational dynamics by interrupting NTD dimerization through steric interference rather than allosteric mechanisms (Schopf *et al*, 2017), and thus presents as a suitable negative control. With the exception of K314 and W320, PRS does not accentuate the HOP binding residues in the FO-ATP, confirming that cofactor binding residues identified by PRS are likely allosteric modulators of conformational dynamics.

## Conclusion

To date, numerous biochemical and computational studies have made considerable advances towards understanding how Hsp90 modulates its complex conformational cycle. In this study, we have assessed the current opinions regarding Hsp90’s conformational dynamics, with respect to the human Hsp90α isoform, in a comprehensive analysis of full length homology models of the human Hsp90α, using all-atom MD simulations coupled with in depth DRN analysis and PRS techniques.

Within the limitations of all-atom MD simulations, our results are in agreement with previous studies describing a differential effect of bound nucleotide on conformational dynamics in which ATP drives NTD dimerization and the closing transition, while the ADP / apo complexes favour the open Hsp90 dimer (Ali *et al*, 2006; Shiau *et al*, 2006; Dollins *et al*, 2007; Krukenberg *et al*, 2011; Li & Buchner, 2013). Regardless of the conformation, bound ATP was found to ‘tense’ the dimer complex in favour of the closed conformation, while ADP appeared to ‘relax’ the protomers affording a greater degree of flexibility. Bound ADP in the partially open complex increased protomer flexibility, resulting in global conformational rearrangements representing the opening transition and the fully open client loading conformation (Shiau *et al*, 2006; Dollins *et al*, 2007; Southworth & Agard, 2011). Analysis of the change in residue reachability over time confirmed the ‘tensing’ and ‘relaxing’ effect of ATP and ADP respectively, with notable reduction in average path length in the former, and variable increased path lengths in the latter. Locations in the protein recording large changes in reachability corresponded to key functional elements such as ß-strand_1_, helix_1_, ATP-lid, and catalytic loop, implicating these regions in steering conformational dynamics. Betweenness centrality analysis provided a measure for distinguishing sites in the protein responsible for intra-protein communication. These sites were found to correlate closely with known functional sites, demonstrating how BC analysis may be used to identify functional sites on a protein.

PRS was used to probe both the closed and open conformation complexes, to assess each residue’s allosteric potential to affect conformational changes. Analysis of the average response of the whole protein to external force perturbation revealed an inter-protomer allosteric coupling between the NTD and CTD of the closed complexes, in which perturbations arriving at the terminal end of one protomer are felt or ‘sensed’ at the opposite terminal of the other protomer. This observation is in agreement with a recent study demonstrating efficient communication between the two domains, leading to the discovery of a CTD allosteric drug target site (Morra *et al*, 2009; Retzlaff *et al*, 2009; Verkhivker, 2014; Morra *et al*, 2010). In a similar manner, analysis of the open complexes demonstrated allosteric coupling between the NTDs of each protomer, the apo complex showing particular sensitivity to perturbations at the NTDs, and an increased sensitivity of the ATP complex over the ADP complex, suggesting ATP driven allosteric activation of this complex. It is well documented that regulation of Hsp90’s conformational cycle is largely impacted by various co-chaperone interactions (Schopf *et al*, 2017; Krukenberg *et al*, 2011; Flynn *et al*, 2015). The perturbations utilized in PRS provide a novel way of introducing external influences on select residues that could be likened to the natural forces involved in co-factor and client binding. PRS analysis of the closed complexes revealed no single residue perturbations capable of selecting a coordinate change towards the open conformational state. It is thus likely that the opening transition occurs spontaneously without the aid of external binding partners. Interestingly, the highest *C*_*i*_ residues selected by PRS for the closed complexes include residues that make up the primary NTD dimerization site, including ß-strand_1_, helix_1_, ATP-lid, and M-domain catalytic loop, elements that have been previously reported to form an important NTD/M-domain hinge implicated in conformational dynamics (Morra *et al*, 2012). Unlike the closed complexes, PRS analysis of the open conformation complexes revealed several sites in the NTD and M-domain capable of selecting a conformational change towards the closed form. Most of the residues at these sites also correlate to known functional sites, while several map to regions on the surface of the protein believed to be co-chaperone binding sites (Schopf *et al*, 2017) (**Figure 13**). In particular, PRS accentuates a number of residues in the ß-strand_1_, helix_1_, ATP-lid, and catalytic loop regions of the FO-ATP complex. These residues correspond with sites implicated in Aha1 binding interactions, suggesting that perturbations at these sites may arrive naturally through co-chaperone binding interactions. This hypothesis is in keeping with recent reports implicating Aha1 in early ATP-lid mobilization (Schulze *et al*, 2016), acceleration of ATPase activation (Panaretou *et al*, 2002), and allosteric modulation of Hsp90’s conformational dynamics (Blacklock *et al*, 2013). The accentuation of these residues in the FO-ATP but not the FO-ADP complex, reinforces the hypothesis that ATP allosterically activates or primes the open complex for conformational transition towards the closed state.

**Figure 13:**
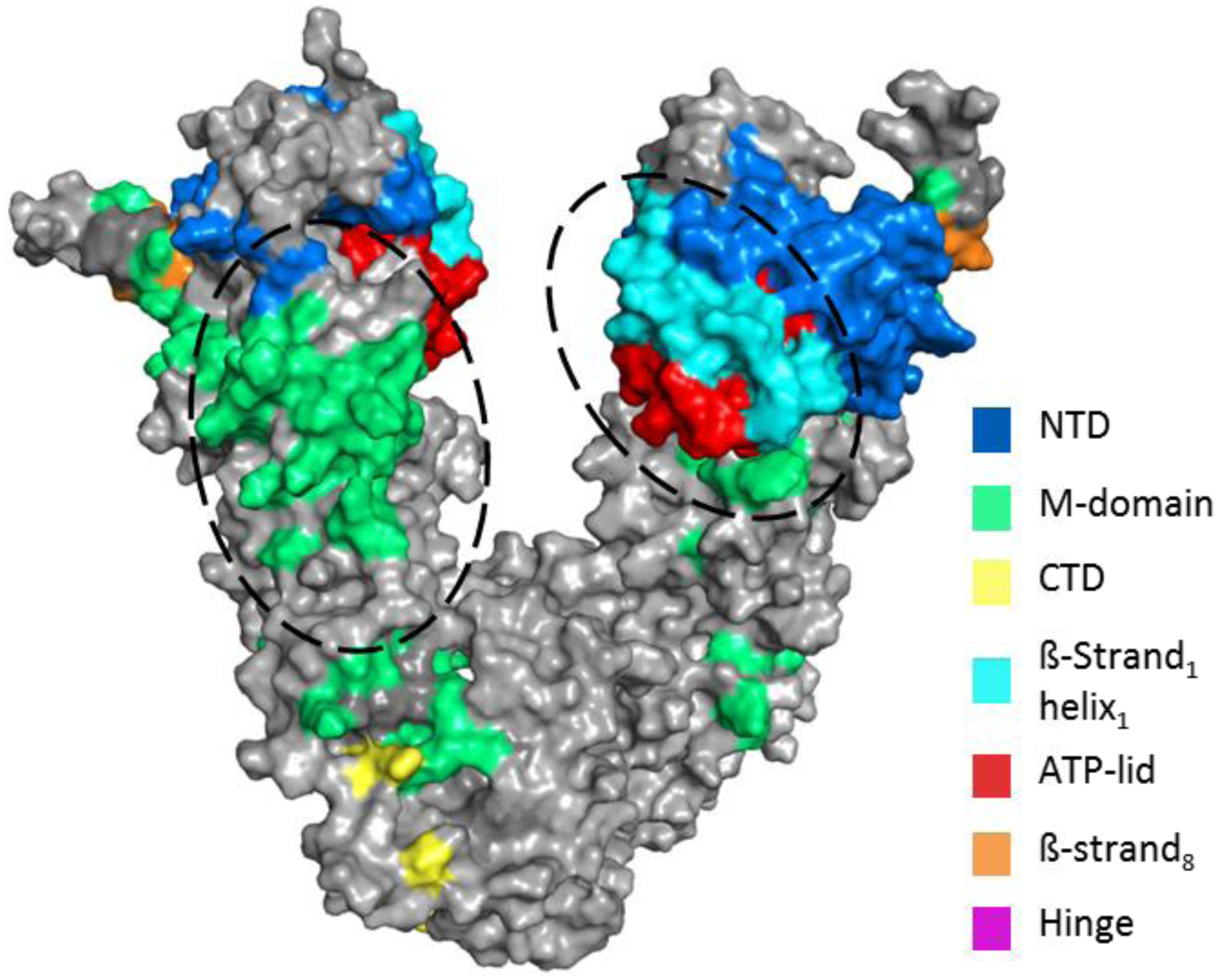
Surface representation of Hsp90 depicting PRS hot spots capable of selecting a conformational change towards the closed complex. Ellipses denote regions that overlap with known co-factor binding sites.

In summary, this study demonstrated how all atom MD simulations coupled with dynamic network analysis and PRS can be used to gain further insights regarding allosteric regulation of conformational dynamics in large complex proteins such as human Hsp90_α_. Collectively, these methods provide a computationally feasible platform to determine functional residues implicated in modulating conformational dynamics, offering routes for novel rational experimental investigations on chaperone function and allosteric regulation.

## Acknowledgements

The authors thank the Centre for High Performance Computing (CHPC), South Africa, for computing time. This work is supported by the National Research Foundation (NRF) South Africa (Grant Number 93690) and the Scientific and Technological Research Council of Turkey (Grant Number 116F229). The content of this publication is solely the responsibility of the authors and does not necessarily represent the official views of the funders.

## Author contributions

ÖTB conceived the project; DLP did all the calculations, most of the analysis and wrote the initial draft of the manuscript; CA and ÖTB contributed to data analysis and revision of the manuscript.

## Supporting information

**SD 1:** Comparison of best scoring homology models before and after energy minimization

**SD 2:** Final closed conformation structures after 100 ns MD simulations for ATP (red); ADP (blue); and apo (green), showing original structure in grey ribbons.

**SD 3:** RMSF plots of closed conformation complexes. Protomer 1 is coloured by NTD configuration (ATP – red, ADP – blue, apo – green), and protomer 2 is in black.

**SD 4:** (A) Final structures of the partially open complexes after 200 ns MD simulations, showing how the ADP structure opens compared to the ATP and apo complexes. (B) Bacterial HtpG structure (PDB 2IOQ) (cyan) superimposed on the 200 ns ADP complex structure (grey), showing the accuracy of the fully open structure.

**SD 5:** RMSF plots of the partially open complexes. The two protomers are labelled A and B.

**SD 6:** (A) RMSD and (B) NTD inter-protomer distance plots for the fully open complex

**SD 7:** RMSF plots for the fully open complexes. Protomer 1 is coloured by NTD configuration (ATP – red, ADP – blue, apo – green), and protomer 2 is in black.

**SD 8:** Summary list of residues accentuated by PRS for the closed (*C*_*i*_ > 0.50) and open (*C*_*i*_ > 0.75) complexes

